# damidBind: an R/Bioconductor package for differential DamID analysis and data exploration

**DOI:** 10.64898/2026.01.06.698059

**Authors:** Owen J. Marshall

## Abstract

**Summary:** DamID, and its cell-type specific adaptations, including Targeted DamID (TaDa) and Chromatin Accessibility TaDa (CATaDa), are now widely-adopted as techniques for the genome-wide profiling of DNA binding proteins. Despite this popularity, no dedicated software solution exists for identifying differentially bound or accessible loci, or differentially transcribed genes, between cell types using DamID. The R/Bioconductor package damidBind provides these functions, allowing an end-user to move from processed binding profiles to identifying differentially-bound loci in a reproducible, statistically appropriate and straightforward workflow.

**Availability and Implementation:** damidBind is an open-source R/Bioconductor package and freely available from Bioconductor at https://bioconductor.org/packages/-damidBind/, and from GitHub at https://github.com/marshall-lab/damidBind. It is released under the GPLv3 licence.

**Contact:** Owen Marshall

## 1 Introduction

DamID-seq profiles genome-wide protein–DNA association by tethering DNA adenine methylase to a DNA-binding or chromatin-associated protein, producing local GATC methylation that can be detected by methylation-sensitive digestion and sequencing(van Steensel and Henikoff, 2000; van Steensel *et al*., 2001; Marshall *et al*., 2016). Cell-type-specific adaptations such as TaDa (Southall *et al*., 2013; Marshall *et al*., 2016; Cheetham *et al*., 2018; Tosti *et al*., 2018) extend this approach to intact tissues without cell sorting or antibodies.

As well as profiling transcription factors and chromatin proteins, DamID-seq can be used to profile the occupancy of RNA polymerase over gene bodies (Southall *et al*., 2013). This readout provides a proxy for gene transcription, and has been successfully used for transcriptional profiling of different cell types (e.g. Southall *et al*., 2013; Marshall and Brand, 2017; Otsuki and Brand, 2018; Doupé *et al*., 2018; Gervais *et al*., 2019).

As Dam has a strong affinity for open chromatin, the DamID technique always obtains a Damonly control profile alongside the Dam-fusion profile, and the final binding profile is a normalised log_2_ ratio of binding. The Dam-only profile can be used as a direct measure of cell-type-specific chromatin accessibility, providing a readout (Chromatin Accessibility TaDa, or CATaDa) similar to ATAC-seq (Aughey *et al*., 2018).

Multiple software tools exist for processing raw DamID-seq NGS reads to log_2_ ratio binding profiles and peak calls, including `damidseq_pipeline` (Marshall and Brand, 2015), `Daim` (Tosti *et al*., 2018) and `Damsel` (Page *et al*., 2024). However, because DamID-seq produces log_2_ ratios over unevenly-spaced GATC fragments, standard ChIP-seq and RNA-seq differential analysis tools are poorly suited to these data.

The R/Bioconductor package `damidBind` addresses this gap, formalising the process of performing differential binding analysis on all forms of DamID-seq datasets. It applies the data curation and underlying statistical analysis appropriate for each DamID application, providing a reproducible, data-appropriate workflow from processed NGS data to biological insights.

## 2 Implementation

### 2.1 Architecture

`damidBind` is implemented as an R package within the Bioconductor (Gentleman *et al*., 2004) ecosystem. Genomic data are stored as `GenomicRanges` (Lawrence *et al*., 2013) objects, linked to genome annotations from `AnnotationHub`. Core statistical tests for differential analysis are handled by `limma` (Smyth, 2005) and `NOISeq` (Tarazona *et al*., 2015), and results are returned as a `DamIDResults` S4 object containing results tables, together with input file and genome annotation metadata. Where applicable, parallel execution is implemented via `BiocParallel` (see Table S2 for benchmarking details).

### 2.2 Input data

The `damidBind` package assumes that the user has processed NGS files to bedGraph profile tracks. For differential protein binding or chromatin accessibility, the package also requires a predetermined set of peak calls for each condition, which can be provided as either external peak files (BED or GFF) or as a `GenomicRanges` R object.

A recommended workflow is NGS alignment and profile generation via `damidseq_pipeline` (Marshall and Brand, 2015), and then peak-calling via either `find_peaks` (Marshall *et al*., 2016) (external, per replicate) or `Damsel` (Page *et al*., 2024) (within R, per condition) (Fig S1; Table S1), before commencing `damidBind` analysis (Fig. 1A).

**Figure 1:**
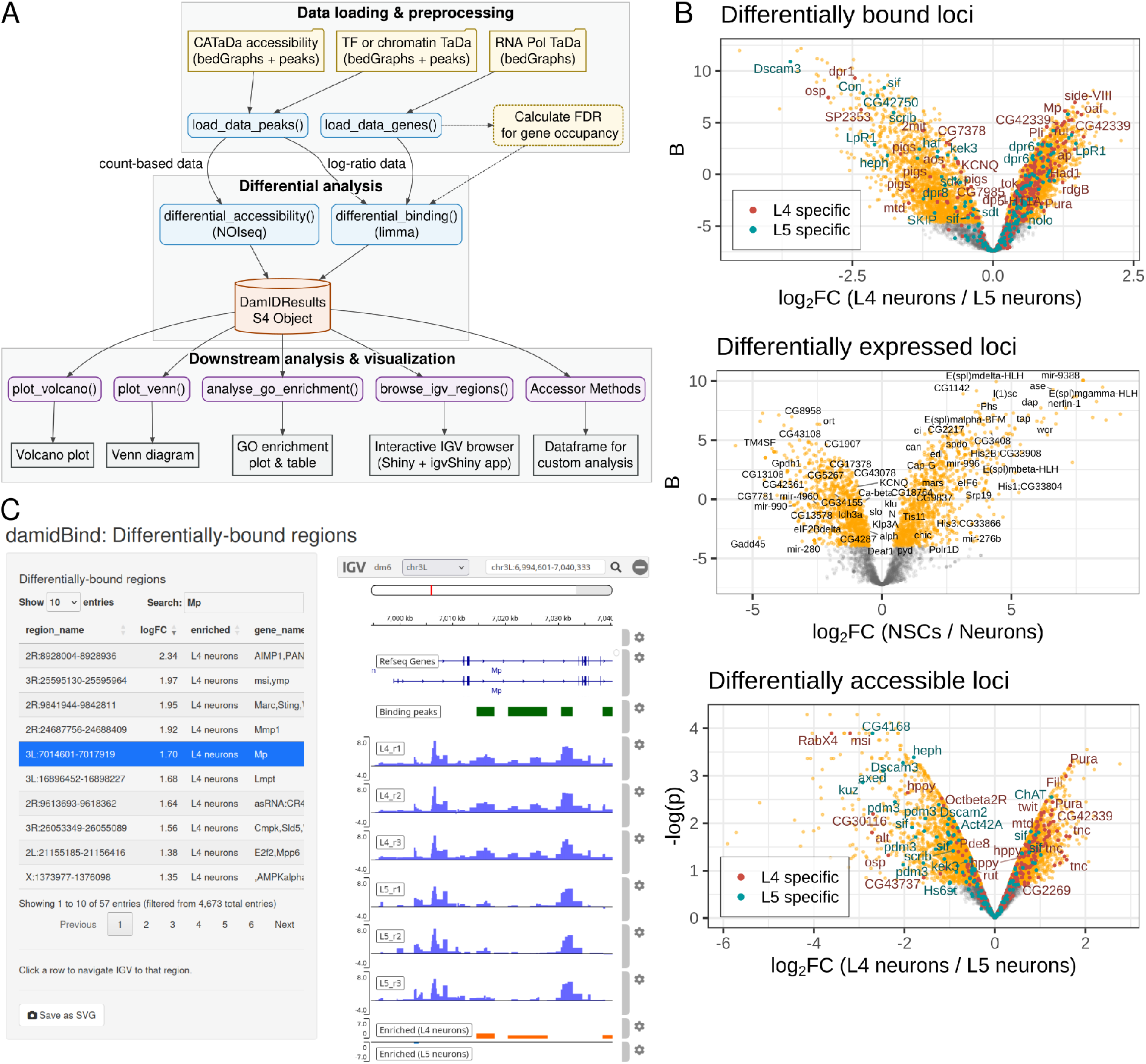
The damidBind package. (A) Schematic overview of the main functions and workflow of damidBind. (B) Example volcano plots of differential binding, expression and accessibility, illustrating different plot labelling options; data are from (Xu *et al*., 2024; Marshall and Brand, 2017, 2015). (C) A screenshot of the Shiny /IGV based interface for interactive locus browsing.

At least three biological replicates per condition are recommended. Analyses with two replicates are supported, but have limited statistical power.

### 2.3 Data handling and normalisation

DamID-seq data are a distinct form of genomic binding/accessibility data which relies on methylated adenines in the sequence GATC. As a result, the underlying resolution of DamID data is the unevenly-spaced distribution of this sequence in the genome. This uneven GATC spacing makes standard count-based differential analysis tools inappropriate for DamID.

At the core of `damidBind`’s analysis is a principle of maximising statistical power by reducing the number of statistical comparisons. For peaks-based analysis pipelines (CATaDa and conventional TaDa), per-replicate or per-condition peak calls are reduced to a set of unified binding regions. For expression analysis, gene body coordinates, as obtained from the most up-to-date genome release (or any historical release) via `AnnotationHub`, are used directly as binding regions.

The variable GATC fragment widths mathematically require spatial weighting to accurately represent locus occupancy. `damidBind` calculates the average, fragment-length-weighted signal for each replicate, ∑_*i*_ *signal*_*i*_ × *width*_*i*_/_*i*_ *width*_*i*_ for all fragments *i* within a locus, where *signal*_*i*_ is either the fragment log_2_ enrichment for DamID binding data, or the reads-per-million count for CATaDa. This method correctly accounts for the variable fragment length structure of loci without being strongly influenced by length-based signal bias (see Supplementary material S.2.1; Fig. S2). The locus-level weighted-means are used in all downstream analysis.

For all workflows, the option to apply uncentered scaling of the input data (default) along with quantile, cyclic LOESS or reads per million (RPM) normalisation prior to locus occupancy calculation is provided (see Supplementary Material S.2.2 for further discussion of data normalisation strategies). Following data loading, `damidBind` provides diagnostic PCA and correlation heatmap plots of the input data, derived from both the underlying raw data and the mean locus occupancy values, allowing the user to check the data quality and condition separation.

### 2.4 Differential analysis workflows

#### 2.4.1 Protein binding

For conventional DamID or TaDa experiments, the `damidBind` workflow is initiated via `load_data_peaks()`. This function imports log_2_(Dam-fusion/Dam-only) binding profiles and associated peak calls, unifies peak regions across all replicates and calculates per-replicate peak occupancy scores. Peaks are also associated with genes in close proximity (within ± 1kb of the gene body by default).

The subsequent analysis with `differential_binding()` uses the empirical Bayes (eBayes) framework of `limma` to identify statistically significant changes in protein binding between two conditions. `limma` is designed to handle heteroscedastic log_2_-ratio data with a mean-variance trend, as is typical in log_2_ ratio DamID binding data, and `damidBind` by default uses the `trend` and `robust` options of `limma`’s eBayes moderation to account for this (Fig S3).

#### 2.4.2 RNA Polymerase occupancy

DamID can be used to profile RNA Polymerase occupancy across gene bodies as a proxy for gene expression. `damidBind` supports this via the `load_data_genes()` function, which calculates replicate occupancy over annotated genes. Differential analysis then proceeds using the same `limma`-based `differential_binding()` function as described above.

For RNA Polymerase datasets, `damidBind` provides the ability to determine the FDR of enriched gene body occupancy per condition, as a proxy for gene expression status. The algorithm used here is a substantially revised version of the approach described in Southall *et al*. (2013), introducing log-linear modelling with weighted natural splines and Jensen’s correction to model heteroscedasticity across fragment counts, and taking advantage of condition replicates to enrich statistical power (see Supplementary Material S.2.5 for algorithm details). The occupancy FDR values are not used directly in differential expression analysis, but can be used to limit downstream analysis and visualisation plots to expressed genes only.

#### 2.4.3 Chromatin accessibility

The Dam-only control sample from a DamID experiment can be used as a proxy for chromatin accessibility, via a method termed CATaDa (Aughey *et al*., 2018). Unlike log_2_ binding ratios, CATaDa data consist of count-derived, non-negative, non-normally-distributed accessibility signals, summarised as fragment-length-weighted averages over peaks. These data are under-dispersed locus-level weighted-average intensities rather than raw counts, and therefore do not naturally satisfy the raw-count assumptions of negative binomial workflows such as `edgeR` (Chen *et al*., 2025) or `DESeq2` (Love *et al*., 2014) (Fig. S4). The package’s `differential_accessibility()` function instead uses `NOISeq` (Tarazona *et al*., 2015) for differential analysis, a package designed to analyse normalised count-derived data non-parametrically, making it robust to this data type (see Supplementary Material S.2.3, Fig. S5 and Table S7 for further discussion and comparison with `edgeR`).

### 2.5 Scope and limitations

At the time of release, `damidBind` only handles pairwise comparisons between datasets. However, the underlying `limma` contrasts framework allows more complex analyses, and these are planned for future releases. More complex CATaDa designs would require separate implementation and validation.

While `damidBind` is primarily designed to compare the same Dam-fusion protein across different contexts, the package can potentially be used to compare the binding of two different proteins in the same cell type (e.g. Nguyen *et al*., 2026). However, users should approach such analyses with caution, as different TFs may have markedly different binding profile distributions, making interpretation of differential binding and the assumptions of shared condition-level residual variance less secure.

### 2.6 Visualisation and downstream analysis

`damidBind` provides a number of tools for downstream functional analysis and publication-ready plots and figures. These include proportional Venn diagrams using `biovenn` (Hulsen *et al*., 2008), Gene Ontology enrichment analysis via `clusterProfiler` (Yu *et al*., 2012), customisable volcano plots (Fig. 1B; see also Supplementary Material S.2.4), and a `Shiny` /`igvShiny` interface for browsing significant loci with replicate tracks, peaks and differential results (Fig. 1C). Because a core feature of `damidBind` is the association of bound loci to neighbouring genes, differential binding analysis is immediately linked to biological function in these analyses.

## 3 Application and validation

### 3.1 Differential binding and accessibility in lamina neurons

To assess the differential binding analysis of `damidBind`, the package was applied to a published TaDa dataset profiling the binding of the transcription factor Brain-specific homeobox (Bsh) in two related neuronal subtypes, L4 and L5 (Xu *et al*., 2024). Xu *et al*. (2024) used independent scRNA-seq data to identify marker genes uniquely transcribed in each neuronal subtype, finding that loci with enriched Bsh binding in Notch-ON L4 neurons were significantly associated with L4 specific markers. In contrast, in Notch-OFF L5 neurons, Bsh loci were depleted for L5 specific markers (Xu *et al*., 2024).

Compared against the analysis from (Xu *et al*., 2024), `damidBind` identified almost all differentially bound loci identified in the earlier study (100% of L4 loci; 92% of L5 loci; Fig. S6A, Table S6). The package also found substantial additional genes differentially bound by Bsh in both neuronal subtypes. Importantly, these additional targets showed the same association of loci with subtype-specific markers as the earlier study, with additional L4 loci enriched for L4-specific gene expression (odds ratio 2.33; P < 1.5 × 10^−10^), and L5 loci not linked with L5-specific gene expression (odds ratio 0.46; non-significant) (Fig. S6A, Table S4). These data suggest that the `damidBind` reanalysis identified subtype-specific Bsh targets, consistent with increased sensitivity without obvious loss of biological specificity in this dataset.

A comparison between the original and `damidBind` analysis of the CATaDa samples in this paper was similar, with `damidBind` recovering 100% and 99% of all loci identified as significant in (Xu *et al*., 2024), and recovering additional subtype-specific expressed genes at a similar odds ratio (Fig. S6B, Tables S8, S9).

### 3.2 Differential transcription between neural stem cells and neurons

To assess the ability of `damidBind` to perform differential expression analysis, Targeted DamID RNA Polymerase II data for *Drosophila* larval neural stem cells and adult neurons was compared with an independent `DESeq2` analysis of FACS-sorted bulk RNA-seq data from broadly corresponding cell types and developmental stages (data from Berger *et al*., 2012; Perlegos *et al*., 2024) (Fig. S7).

As expected for an orthogonal comparison between polymerase occupancy and steady-state transcript abundance, concordance was substantial but incomplete, with a logFC correlation of 0.52 across all differentially-expressed genes in the datasets (Fig. S7A,B), a directional concordance of 71% across all significant genes, and 80% for significant genes called by `damidBind`. When limited to genes with |logFC| > 2, this rose to >80% for both methods (Table S10). Gene Ontology analysis showed that genes significant in both datasets were enriched for expected neural development/function categories, whereas RNA-seq-specific abundance changes were enriched for metabolic, splicing and ribosome biogenesis terms rather than neural development (Fig. S7C). These differences may reflect post-transcriptional regulation, transcript half-life and driver/developmental-stage differences between the datasets.

Differential analysis of DamID-seq RNA Polymerase occupancy therefore recovers biologically meaningful, cell-identity-associated transcriptional differences, while remaining distinct from steady-state RNA abundance.

## Conclusions

`damidBind` provides a standardised, comprehensive differential analysis package for DamID-seq data, encompassing all three major uses of the technique, with diagnostic plots and data visualisation functions that allow researchers to quickly identify key loci for downstream validation and further investigation. The package comes with a comprehensive vignette, and is freely available through the official Bioconductor software repository.

## Software availability

`damidBind` is available from Bioconductor at https://bioconductor.org/packages/damid-Bind/, with DOI 10.18129/B9.bioc.damidBind. The package source code is also available from GitHub at https://github.com/marshall-lab/damidBind. The archived package version used for the analyses in this manuscript, damidBind v1.1.1, is available from Zenodo at https://doi.org/10.5281/zenodo.20758869.

## Acknowledgements

I thank Jake Newland and Caroline Delandre for helpful comments on the manuscript, ad Andres Wokaty and the Bioconductor community for code review. This work was supported by a National Health and Medical Research Council (NHMRC, Australia) Ideas grant [APP2029752].

## Supplementary material

### S.1 Supplementary methods

#### S.1.1 Data sources

Binding data (including CATaDa data) for Bsh (Xu *et al*., 2024) was obtained from GEO accession number GSE247239. RNA Polymerase occupancy data (Marshall and Brand, 2017) was obtained from GEO accession numbers GSE77860 and GSE69184, processed with `damidseq_pipeline` v1.6.3 with default settings. Bulk RNA-seq data used in this study was obtained from GEO accession numbers GSE235989 (Adult 6D; Elav-GAL4 UAS-mCD8GFP control samples) (Perlegos *et al*., 2024), and GSE38764 (third instar larval brain, wild-type neuroblasts) (Berger *et al*., 2012).

The DamID-seq processed data and peak calls are provided in a Zenodo dataset at doi.org/10.5281/zenodo.16649477. This processed dataset consists of log_2_-ratio binding profiles, CATaDa RPM-normalised bedgraphs, alongside associated peak calls for the Bsh binding and CATaDa data.

#### S.1.2 DamID-seq data processing

DamID-seq data was processed with `damidseq_pipeline` v1.6.3 with either default settings (TF binding and RNA polymerase occupancy) or with the `--catata` flag (Dam-only samples for CATaDa). Peak calling was carried out using `find_peaks` v1.2 with default settings.

#### S.1.3 damidBind analysis

Differential binding /expression analysis was performed using `damidBind` v1.1.1 (Zenodo DOI: https://doi.org/10.5281/zenodo.20758869). Quantile normalisation was used as the default normalisation method unless otherwise specified; *Drosophila melanogaster* BDGP6.46 /Ensembl 113 (AH119285) was used as the genome annotation from `AnnotationHub`. All other settings used package defaults (including a ± 1kb window for gene associations with loci, q=0.8 as the NOISeq significance threshold for CATaDa analysis, and FDR<0.05 to filter for significant gene occupancy for RNA Polymerase II differential transcription analysis).

#### S.1.4 RNA-seq data processing and analysis

Raw FASTQ RNA-seq reads were aligned using `kallisto` v0.46.1, against the *D. melanogaster* BDGP 6.46 release 113, with `--genomebam -t 14` runtime flags. Transcript abundance `tsv` files were imported into R via `tximport`, analysed using `DESeq2`, and the results processed with `lfcShrink` using `apeglm` (Zhu *et al*., 2019).

#### S.1.5 Additional analysis of CATaDa data

To assess the performance of `NOISeq` on CATaDa data, results were compared against the `edgeR` (v4.4.2) quasi-likelihood (QL) framework. Input data consisted of the same lamina neuron fragment-length-weighted RPM scores (summarised over peaks) used in the `damidBind` analysis. These scores were treated as fractional counts within `edgeR`; library size normalisation was performed using the trimmed mean of M-values (TMM) method via `calcNormFactors`, and common, trended, and tagwise dispersions were estimated using `estimateDisp`. Differential accessibility was determined using the `glmQLFit` and `glmQLFTest` functions, with significant loci defined at an FDR < 0.05.

Empirical null-call behaviour was evaluated using a permuted null test on all nine unique non-biological 3-versus-3 partitions (treating complementary partitions as equivalent and excluding the true L4-versus-L5 split, such that each group contained a mixture of biological conditions). Differential analysis was then performed on these pseudo-conditions using both `NOISeq` and `edgeR`.

#### S.1.6 Benchmarking

The main data processing steps for `damidBind` were benchmarked via the `bench` R package. Benchmarking was performed on both the RNA Polymerase II and Bsh TF binding datasets used in this study, using pre-loaded binding profile `GRanges` objects to minimise the contribution of disk I/O. The results are summarised in Table S2, for single core execution vs 15 and 30 cores. Benchmarks were performed on an AMD EPYC Milano server with 32 cores and 128 Gb RAM, running Ubuntu 22.04 and R 4.4.0.

#### S.1.7 Additional data analysis

All additional analysis was performed using R (R_Development_Core_Team, 2011) v4.4.0. Flowchart diagrams were generated using `DiagrammeR`. Composite plots were generated using `Patchwork`.

#### S.1.8 Figure analysis code

The code and data to generate all main and supplementary analysis figures is available at GitHub at https://github.com/marshall-lab/damidBind_manuscript_figures, with an archived snapshot at Zenodo, DOI: https://doi.org/10.5281/zenodo.20758895.

### S.2 Supplementary methodological analyses

#### S.2.1 Effect of fragment-length-weighted averaging

In order to investigate the effects of fragment-length-weighted averaging of DamID-seq data, the relationship between log_2_ ratio signal and GATC fragment length was examined. The observed signal of any given fragment *i* can be considered as

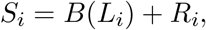

where *B*(*L*_*i*_) is any systematic, fragment-length-dependent bias, and *R*_*i*_ is the residual signal, comprising true biological binding, stochastic technical noise, and any length-independent technical variation. For a genomic locus composed of multiple fragments, the signed discrepancy between the fragment-length-weighted mean and the simple mean is therefore

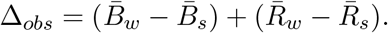

and the corresponding signed discrepancy expected from fragment-length bias alone is

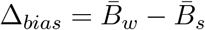

Fragment-length bias was modelled for each sample by excluding all fragments overlapping a binding peak, which were considered to contain true signal for which *R*_*i*_ » *B*(*L*_*i*_). A generalised additive model was then fitted genome-wide to the remaining background fragments:

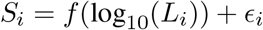

where *f* was estimated using a cubic regression spline. Using this model, the expected length-associated signal 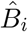 was predicted for all fragments within tested loci. Because the background set may still contain weak or undetected biological signal, this should provide a conservative upper estimate of fragment-length bias.

For each tested locus containing multiple fragments, the observed absolute discrepancy between the fragment-length-weighted mean and the simple mean was calculated as

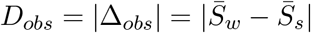

The corresponding absolute discrepancy expected from the fitted fragment-length bias model alone was calculated as

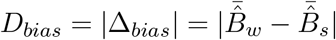

These quantities were compared across tested loci and plotted against within-locus fragment-length heterogeneity, measured as the coefficient of variation of fragment widths (CV).

The DamID-seq background signal was not fully independent of length (Fig. S2A), indicating a weak sample-specific fragment-length bias. However, the observed absolute discrepancy between simple and fragment-length-weighted means substantially exceeded the discrepancy expected from this bias alone. Across the six samples in the Bsh dataset used in this study, the geometric mean ratio of median absolute discrepancies, median(*D*_*obs*_)/median(*D*_*bias*_), was 2.91-fold (95% t-based CI for the population geometric mean: [1.78–4.77]; one-sided paired Wilcoxon signed-rank tests within samples, all *P <* 2 × 10^−16^; Fig. S2B, Table S3).

These results indicate that the effect of fragment-length weighting is not primarily explained by systematic fragment-length bias. Instead, the larger observed discrepancy is consistent with the weighted mean better representing spatial occupancy across uneven GATC fragments.

#### S.2.2 Comparison of differing input normalisation strategies

Lineage-specific factors can exhibit biological asymmetry in binding. One example is Bsh, which in L4 neurons binds at a Notch-dependent open chromatin landscape that is not present in L5 neurons (Xu *et al*., 2024). In order to understand the impact of different normalisation strategies, a simple zero-preserving scaling approach (via R’s internal `scale(center = FALSE, scale = TRUE)`) was compared with cyclic LOESS (the `cyclicloess` method from `limma`) and quantile normalisation. Results were assessed using both subtype-specific scRNA-seq expression markers as a ground truth.

The scaled but otherwise unnormalised dataset showed strong bimodality in the t-statistic distribution, suggesting either unnormalised technical structure, strong biological asymmetry, or both. Quantile normalisation strongly reduced this bimodality, whereas LOESS removed bimodality entirely (Fig S3). However, intersecting the differentially-bound loci from these analyses suggested that LOESS over-corrected the asymmetry of Bsh binding. In L4 neurons, LOESS normalisation appeared to suppress identification of true biological signal, identifying fewer novel binding loci concordant with L4 expression markers (Odds ratio 1.78) than with quantile normalisation (Odds ratio 2.33) (Tables S4, S5). A corollary of this was the identification of over 800 additional L5-enriched loci that lacked any association with L5 subtype markers (Odds ratio 1.12). Conversely, quantile normalisation showed the expected depletion of L5 markers among novel targets (Odds ratio 0.46) (Tables S4, S5).

These results should not be interpreted as a general recommendation of quantile normalisation for all DamID datasets; rather, they illustrate that normalisation choice can affect biological conclusions, and should be carefully assessed against both data diagnostics and known biological priors when performing differential analysis. Both cyclic LOESS and quantile normalisation methods on input data are provided by `damidBind`.

#### S.2.3 Suitability of NOISeq for CATaDa analysis

A mean-variance plot of the CATaDa data from Xu *et al*. (2024) shows that these loci-weighted-average count scores are substantially under-dispersed, violating negative binomial assumptions. While `edgeR` can handle fractional count data, and its quasi-likelihood (QL) framework can potentially handle under-dispersion, the non-parametric analysis of `NOISeq` was hypothesised to be a more appropriate fit for these unusual data.

To determine the suitability of `NOISeq` to handle these data, a direct comparison was performed between `NOISeq` and `edgeR` (using `edgeR`’s quasi-likelihood framework) on the lamina neuron CATaDa data used in this study, using both the true biological data and a permuted null test on all nine unique non-biological 3-versus-3 partitions (treating complementary partitions as equivalent and excluding the true L4-versus-L5 split). `NOISeq` identified 3593 loci in the true biological comparison, but only 26.4 loci on average in the mixed-label null contrasts, with a maximum of 70, corresponding to a mean empirical null-call fraction of 0.7%, and a worst-case fraction of 1.9% (Fig. S5A; Table S7). `edgeR` identified 4368 loci in the true biological contrast and no loci in the mixed-label null contrasts (Fig. S5A; Table S7). These results indicate low false-positive behaviour for both methods under permutation with this example dataset. For the true biological condition, almost all (99.6%) of the loci identified by `NOISeq` were also called by `edgeR` (Fig. S5B-E), providing confidence in the `NOISeq` analysis.

For this representative dataset, `NOISeq` thus appears to represent a conservative choice, and does not appear to unduly inflate Type I errors in the permuted null test. While `edgeR` QL also performed well on the example dataset, the underlying data inputs remain as under-dispersed locus-level weighted average intensities rather than raw counts. These analyses support the use of `NOISeq` as a pragmatic, non-parametric solution for analysing locus-averaged CATaDa intensities.

#### S.2.4 Volcano plot label thinning

Differential analysis typically involves the display of hundreds to thousands of significant loci in volcano plots. Current graphical label packages (e.g. `ggrepel` on thousands of datapoints) within the R ecosystem are limited by total non-overlapping label plot space constraints, meaning that significant point clusters with high density remain unlabelled.

To solve this, `damidBind` uses Algorithm 1 to thin the list of labelled points in dense regions, displaying the thinned list with `ggrepel`. The result is deeper labelling within the significant loci cloud, increasing the number of loci with displayed labels within dense regions of volcano plots, while still preserving labels on isolated points.

#### S.2.5 Gene occupancy FDR estimation

While `damidBind` is primarily concerned with differential analysis, RNA Polymerase DamID, as a proxy for gene expression, allows an additional option of restricting any downstream visualisation to expressed genes only. The gene body occupancy FDR algorithm used for this purpose by `damidBind` is substantially modified from the original described in Southall *et al*. (2013), using a two-tiered regression modelling approach built upon a permutation-based null distribution. The objective is to create a predictive model (Algorithm 2) that can estimate an empirical *p*-value for any given gene’s observed occupancy score and fragment count (Algorithm 3). The per-gene *p-*values from each replicate are grouped by test condition during the

##### Algorithm 1

Sampling points by local isolation

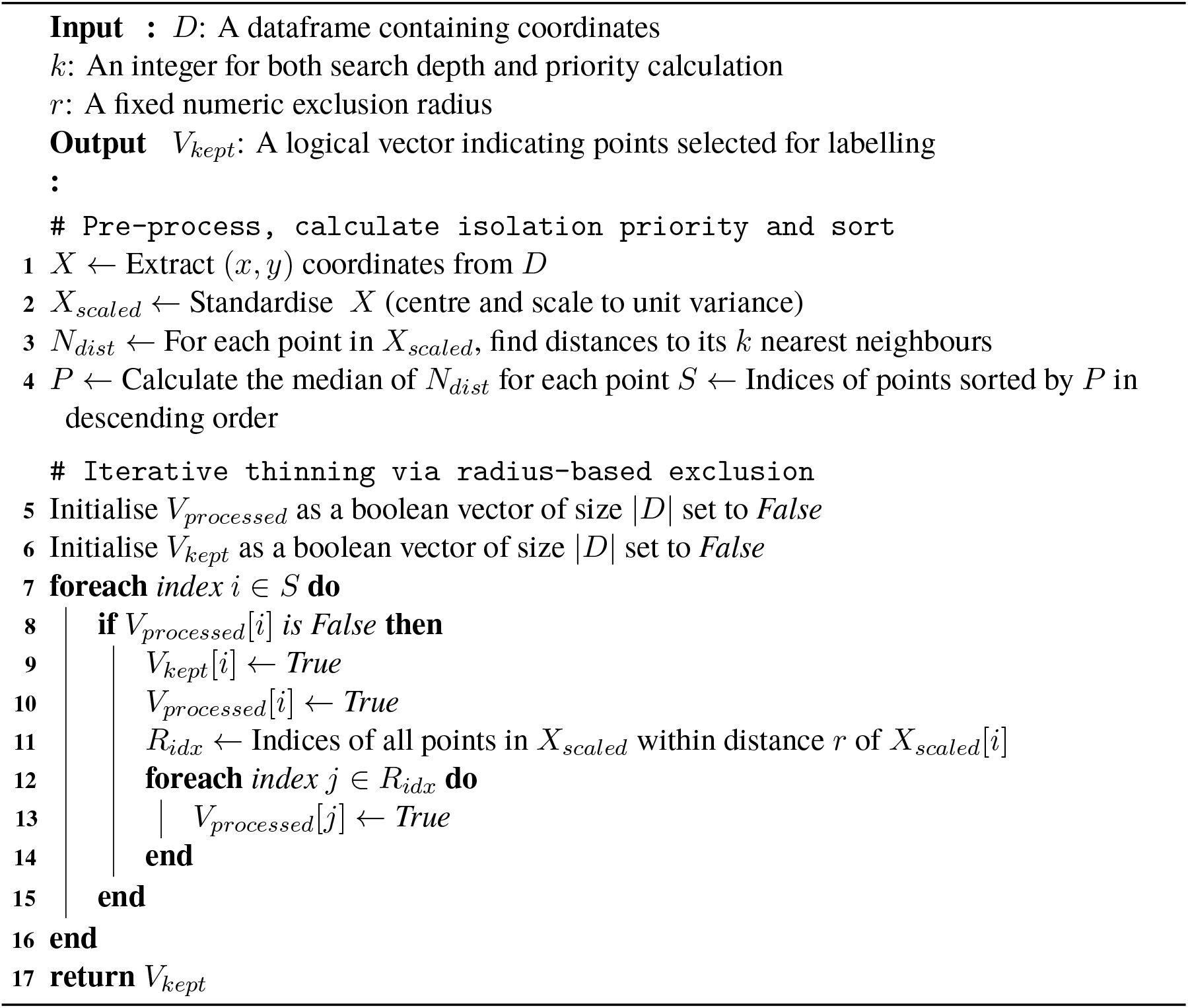

`differential_binding()` call and aggregated using either Fisher’s (Fisher, 1925) or Stouffer’s (Stouffer *et al*., 1949) method, to give a per-condition *p*-value for each gene. Finally, these are adjusted for multiple hypothesis testing using the Benjamini-Hochberg method (Benjamini and Hochberg, 1995) to yield a final per-gene occupancy FDR.

When applied to correctly-normalised RNA Polymerase DamID data, the occupancy FDR serves as a proxy for gene expression (Southall *et al*., 2013), although genes with significant paused RNA Polymerase may also be detected using this approach. The gene occupancy FDR values are not directly used to determine differential gene expression changes, but are used to filter the downstream plots and analysis for expressed genes only.

The major algorithmic changes from Southall *et al*. (2013) in the method implemented in `damidBind` include:

1. The use of fragment-length-weighted mean values for building the null distribution.
2. Generating the underlying null model distributions by sampling with replacement.
3. The use of log-linear models for the Tier 1 regressions.
4. Using natural spline fits for the Tier 2 regressions (slope, intercept, mean squared error (MSE) of residuals) to capture non-linear relationships across the range of fragment counts.
5. The use of weighted least squares (WLS) for Tier 2 slope and intercept models, using the inverse of the Tier 1 standard errors as weights, to account for model heteroscedasticity (Baskerville, 1972).
6. Applying a bias correction factor derived from the predicted MSE when back-transforming from the log-scale, to correct for Jensen’s inequality (Finney, 1941; Baskerville, 1972).
7. Determining a condition-level FDR by combining unadjusted replicate p-values using either Stouffer’s Z-score method or Fisher’s method, followed by Benjamini-Hochberg (BH) adjustment at the condition level.

In order to capture the non-linear relationship between fragment counts and the regression parameters for the Tier 2 regressions, the slope and intercept are modelled as a function of the log-transformed fragment count (Fig. S8A, B). The pseudo-linear residual variance (MSE) is modelled on the linear fragment count (Fig. S8C) to accurately represent the variance profile in low fragment count genes.

##### Algorithm 2

Occupancy FDR: Model training

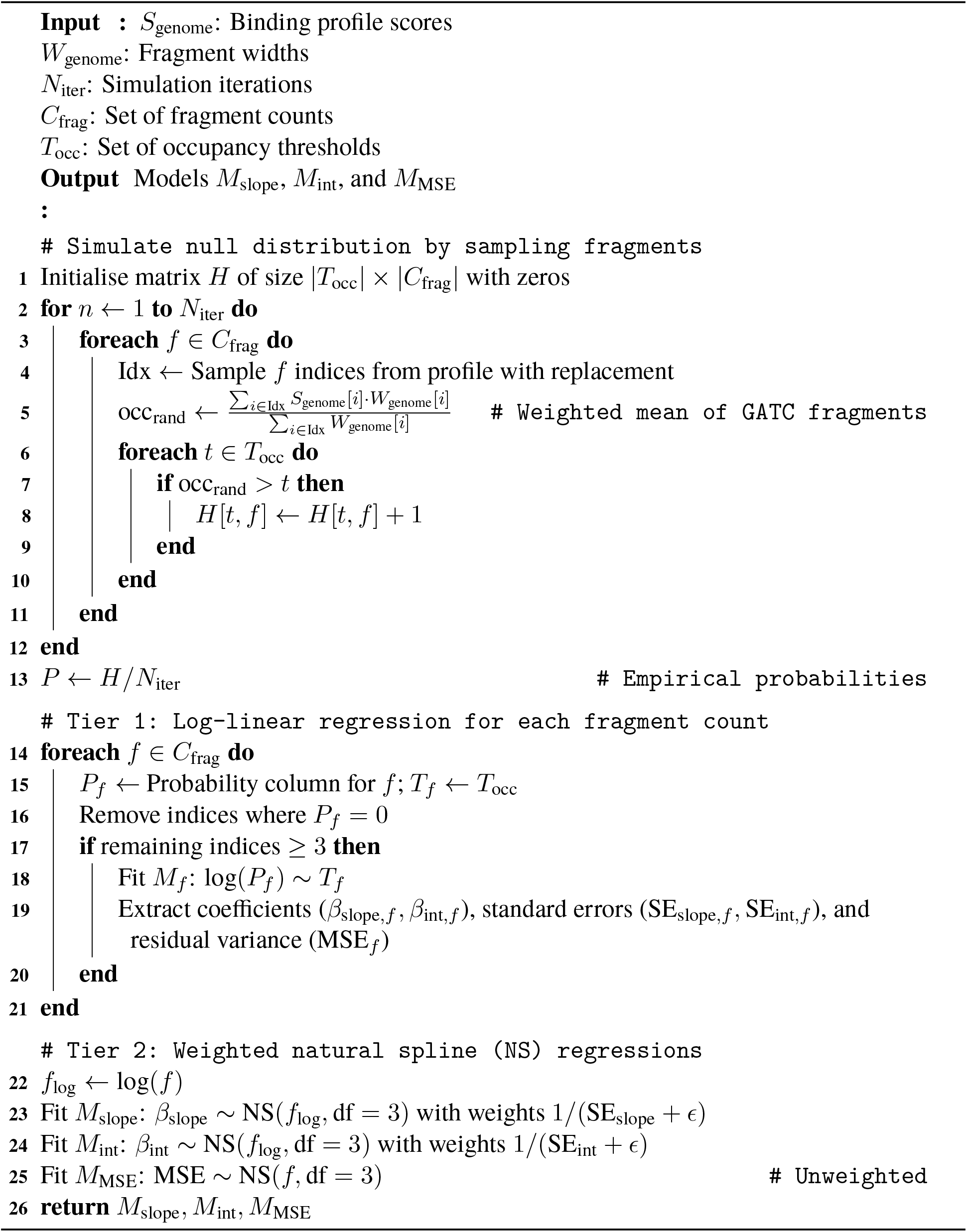

##### Algorithm 3

Occupancy FDR: *p*-value estimates and replicate integration

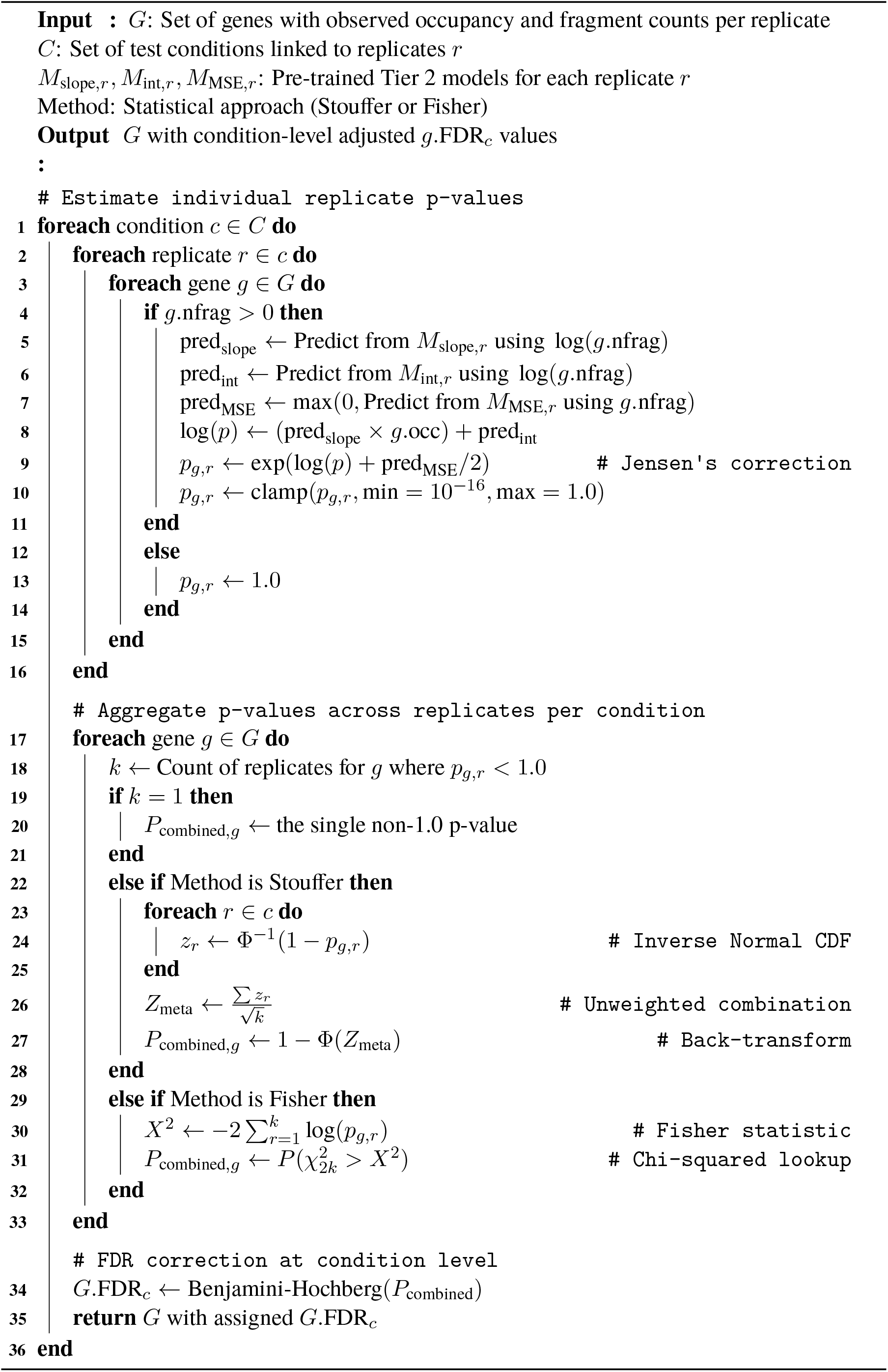

## Supplementary figures

**Figure S1:**
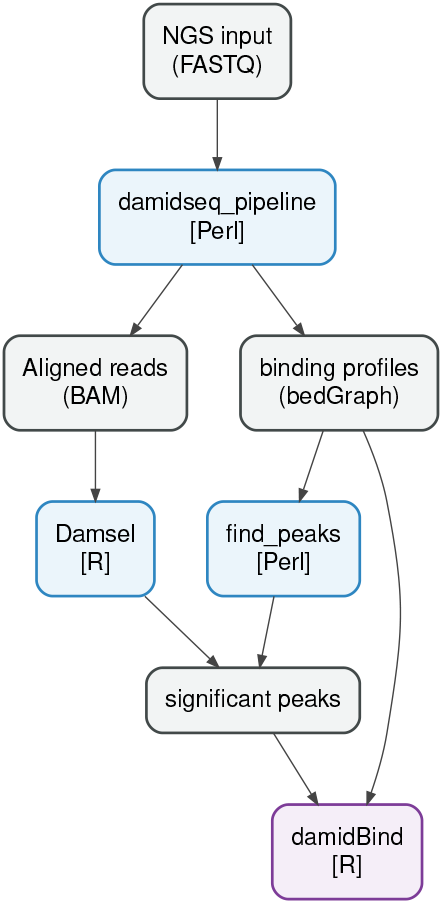
Suggested workflow options for DamID-seq data processing, from raw sequencing reads to `damidBind`.

**Figure S2:**
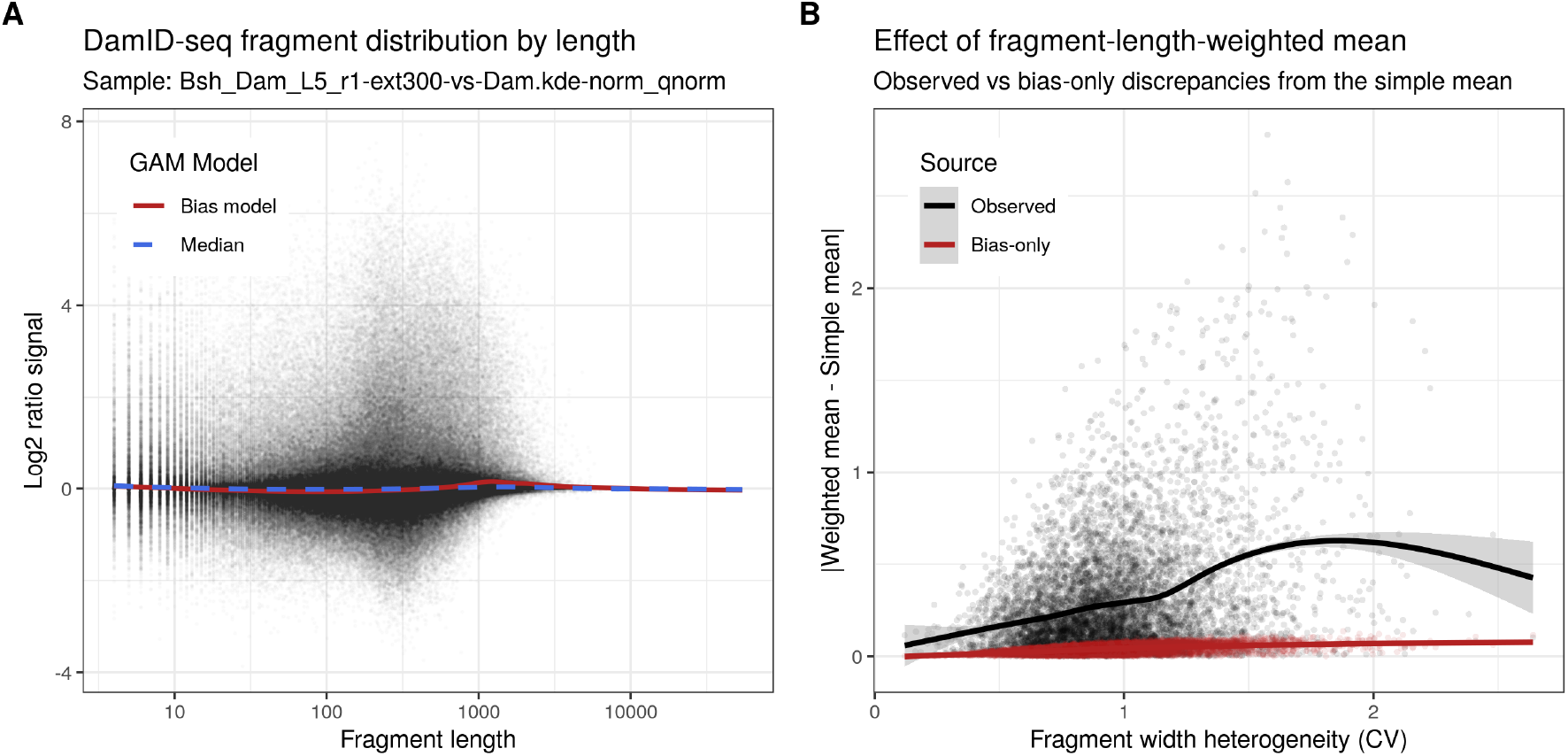
Discrepancies between fragment-length-weighted averaging and simple averaging are not driven by systematic bias. (A) Plots of the underlying fragment length bias over all non-peak GATC fragments, used to fit a null model of systematic fragment-length signal bias (red); a fit of the signal median is also shown. (B) Discrepancy between simple and fragment-length-weighted means, both from the data (Observed, black) and as predicted using the bias-only model (Bias-only, red).

**Figure S3:**
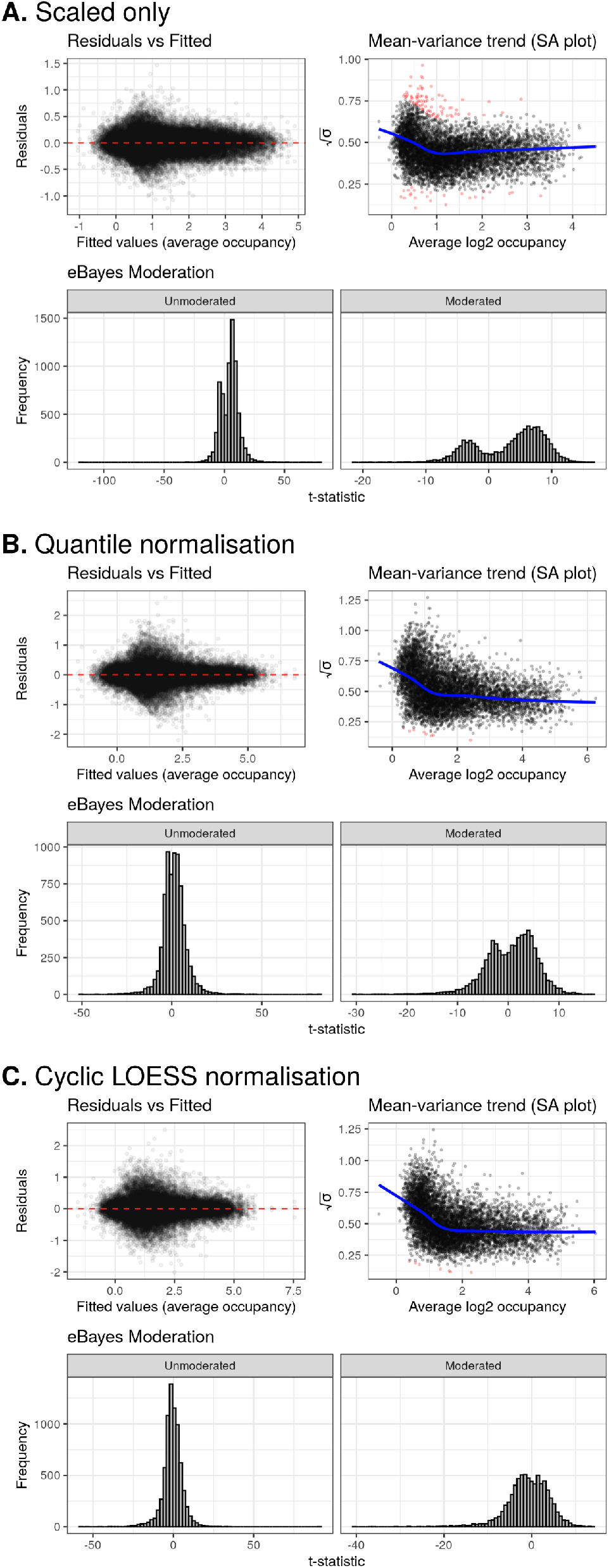
Diagnostic plots for `limma`’s eBayes moderation generated by `damidBind` using Bsh L4/L5 binding dataset with only zero-preserving scaling, with quantile normalisation, and with cyclic LOESS normalisation. Plots showing the heteroscedasticity of the data, the SA plot showing the model fit to the mean-variance of the data (outliers coloured red; fit line in blue), and the effect of eBayes moderation on t-statistic shrinkage.

**Figure S4:**
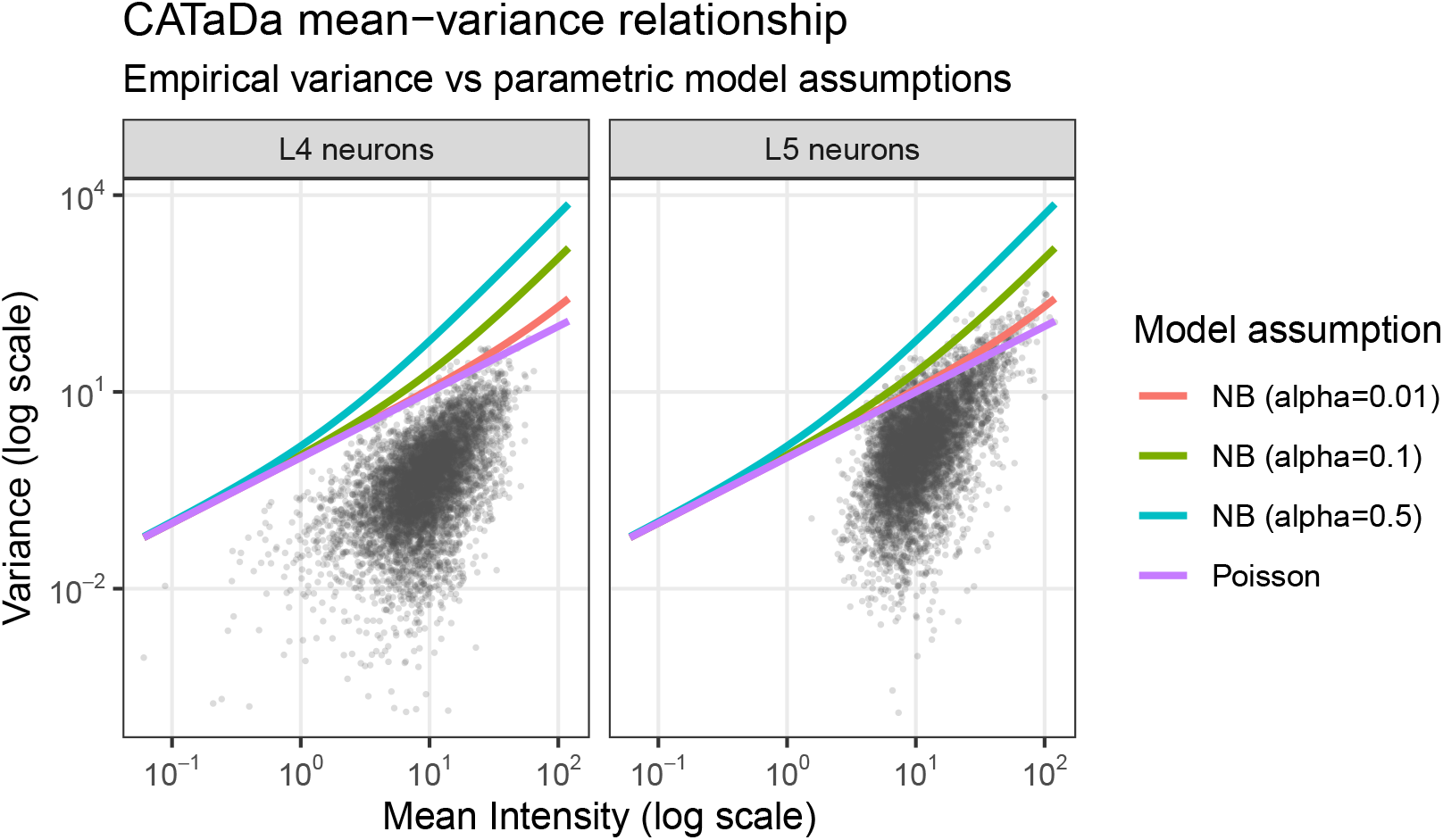
Plots of mean vs variance relationship of CATaDa peak data. Data for L4 and L5 neuronal subtypes (from (Xu *et al*., 2024)) are shown, alongside Poisson (Var = *µ*) and Negative Binomial (Var = *µ* + *αµ*^2^) responses; points represent individual loci. Input data was loaded with `pre_scale=FALSE, norm_method=“none”` (i.e. no normalisation was applied to the dataset). In both cases, the data sit below the model lines and are under-dispersed, violating NB and Poisson assumptions.

**Figure S5:**
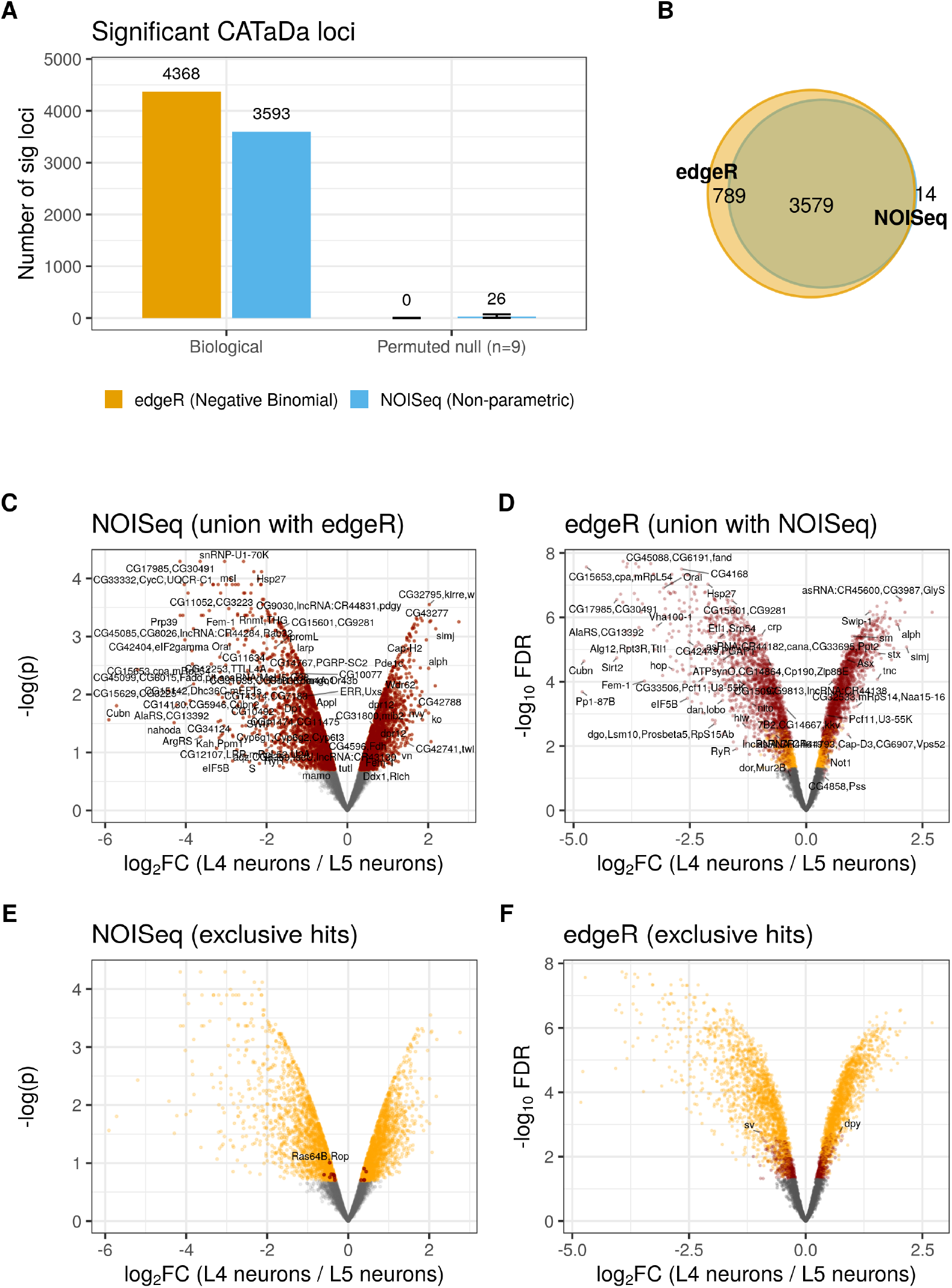
Comparison of `NOISeq` and `edgeR` performance on CATaDa data. (A) Significant hits generated by both methods, for both the biological lamina neuron data from Xu *et al*. (2024) and a permuted null dataset which should yield no hits, to assess FDR performance. (B) Venn diagram of overlapping significant loci called by the two methods on the lamina dataset. (C and D) Significant `NOISeq` and `edgeR` loci, coloured by set overlap in red, other significant loci in orange. (E and F) Significant `NOISeq` and `edgeR` loci, coloured by set uniqueness in red, other significant loci in orange.

**Figure S6:**
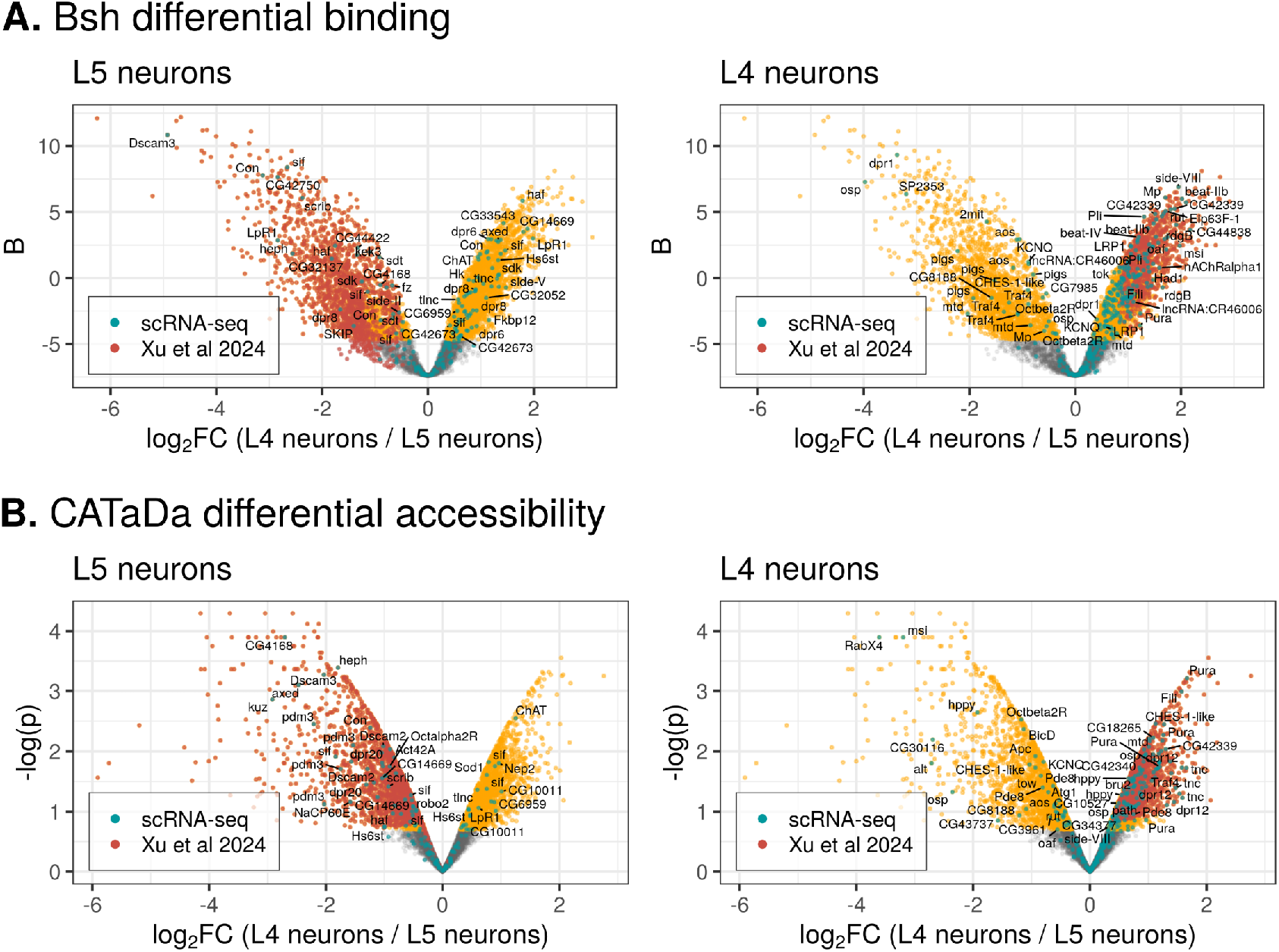
Comparison of `damidBind`’s performance against previously published analysis from Xu *et al*. (2024). (A) Bsh TF differential binding; (B) CATaDa differential accessibility. In all plots, significant loci identified by `damidBind` are coloured orange; significant loci identified by Xu *et al*. (2024) are coloured red; scRNA-seq gene expression markers uniquely expressed in each lineage (data from (Xu *et al*., 2024)) are coloured cyan. See also Table S4. Results using quantile normalisation are shown.

**Figure S7:**
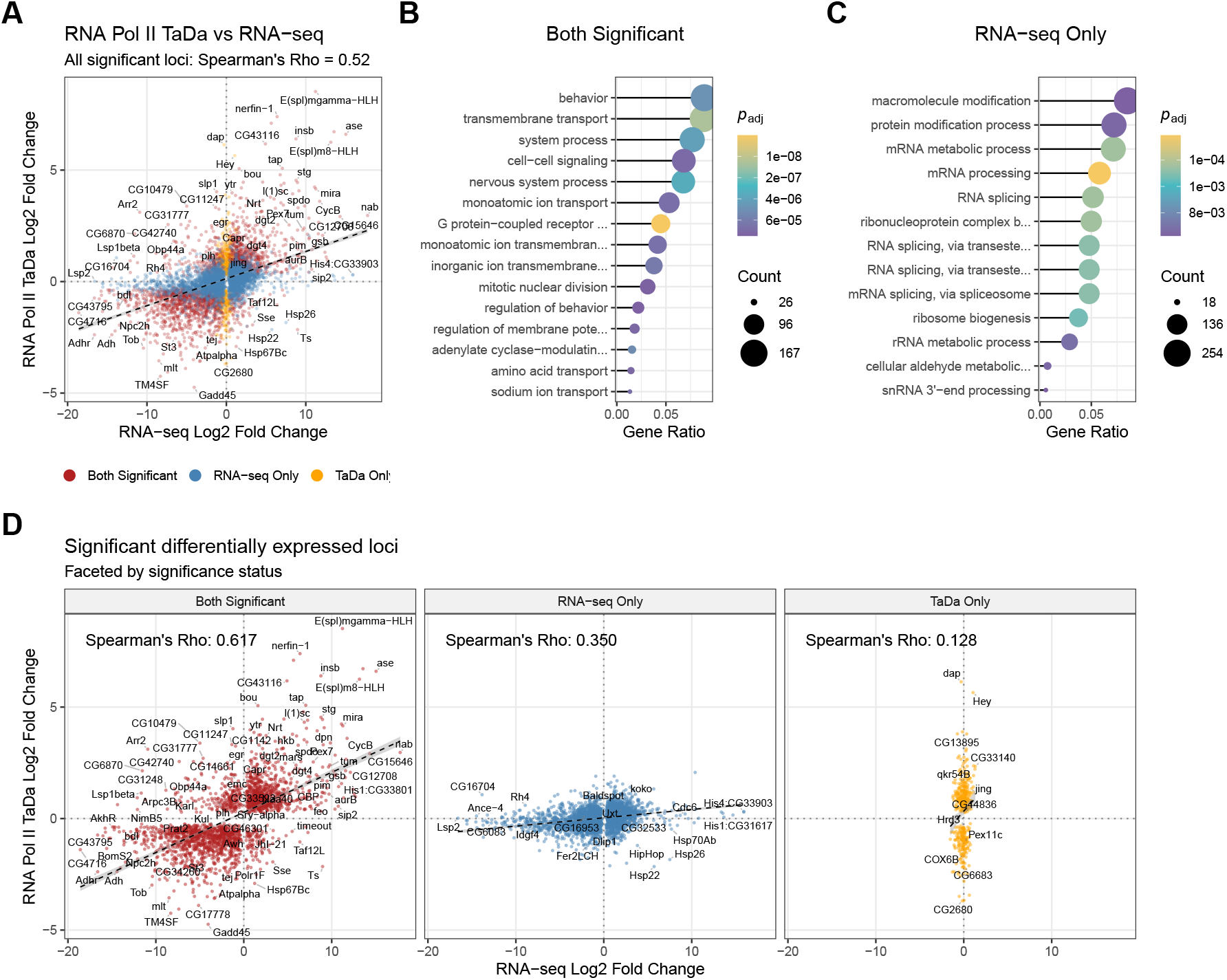
Differential expression compared between TaDa and RNA-seq. In both cases, third instar larval neural stem cells were compared to adult neurons. (A) All significant loci compared; (B) faceted by significance status; (C) Gene Ontology (GO) enrichment terms by significance class (no GO terms were significantly enriched for the TaDa-only significance group). See Supplementary Methods for details.

**Figure S8:**
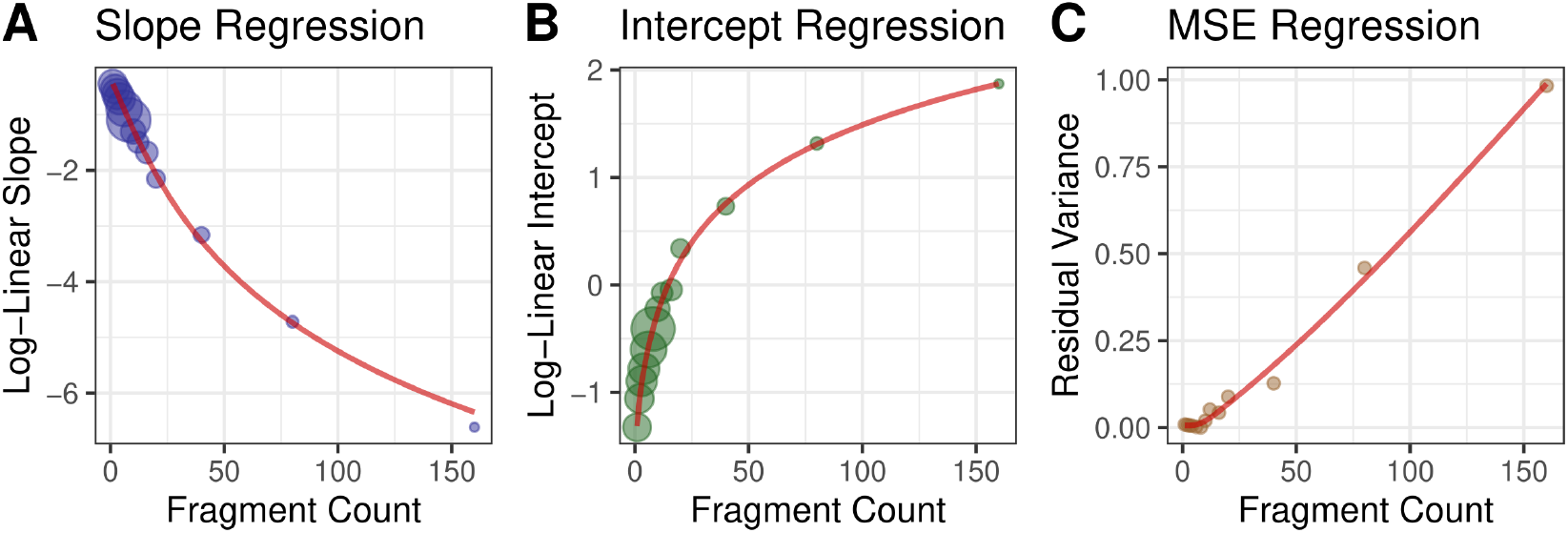
FDR modelling for RNA Polymerase occupancy data. The plots show the Tier 2 regression fits and Tier 1 model heteroscedasticity, for a typical DamID RNA Polymerase dataset. Both the Tier 2 slope (A) and intercept (B) regressions use weighted natural spline fits to fit the underlying data; circle size is proportional to weight. The model mean squared error (MSE) increases with fragment count (C) and is also fitted with an unweighted natural spline to apply the appropriate model bias correction upon back-transformation. Data illustrated is derived from RNA Polymerase (RpII215) occupancy data from GSE77860.

## Supplementary tables

**Table S1:**
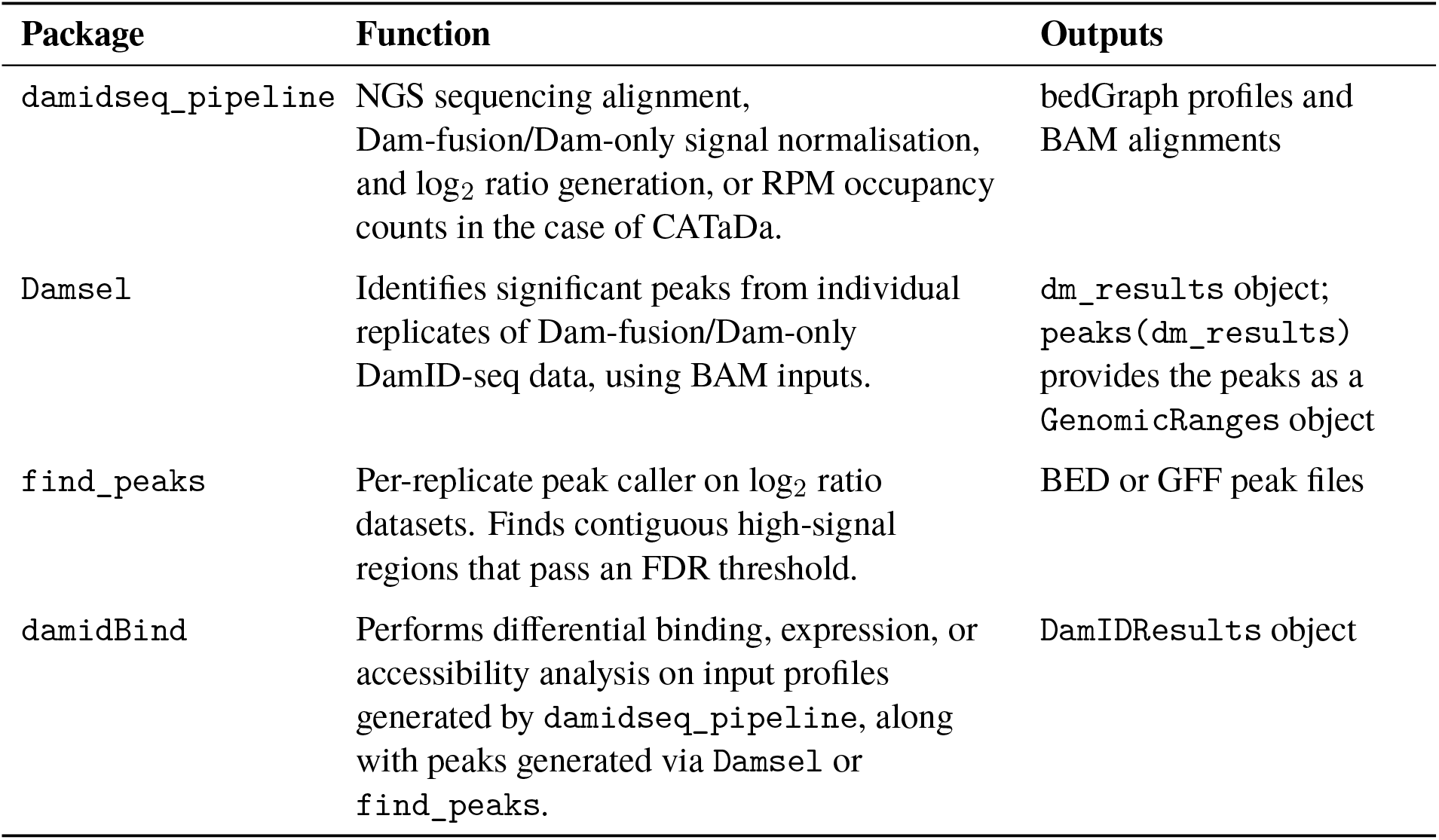
Summary of DamID-seq-related bioinformatics software.

**Table S2:**
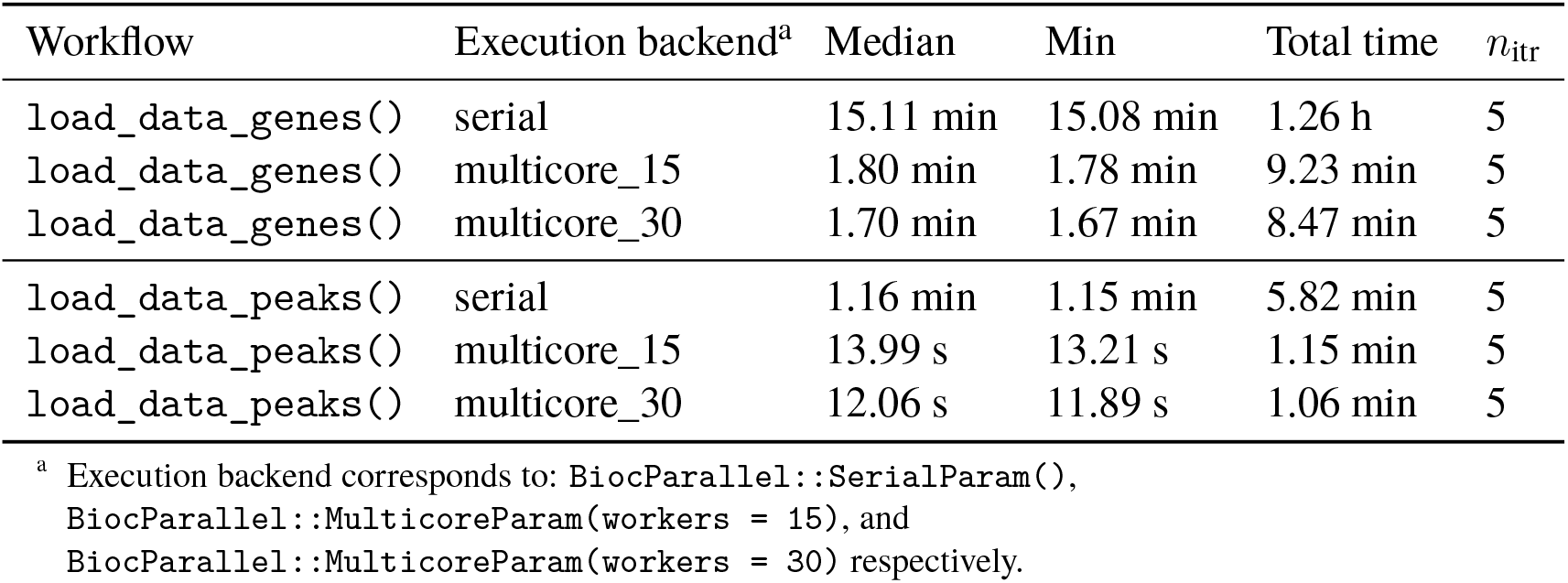
Benchmark results for damidBind workflows using serial and multicore execution. Significantly shorter execution times result from parallel processing, although the benefits of using more than 15 cores is limited.

**Table S3:**
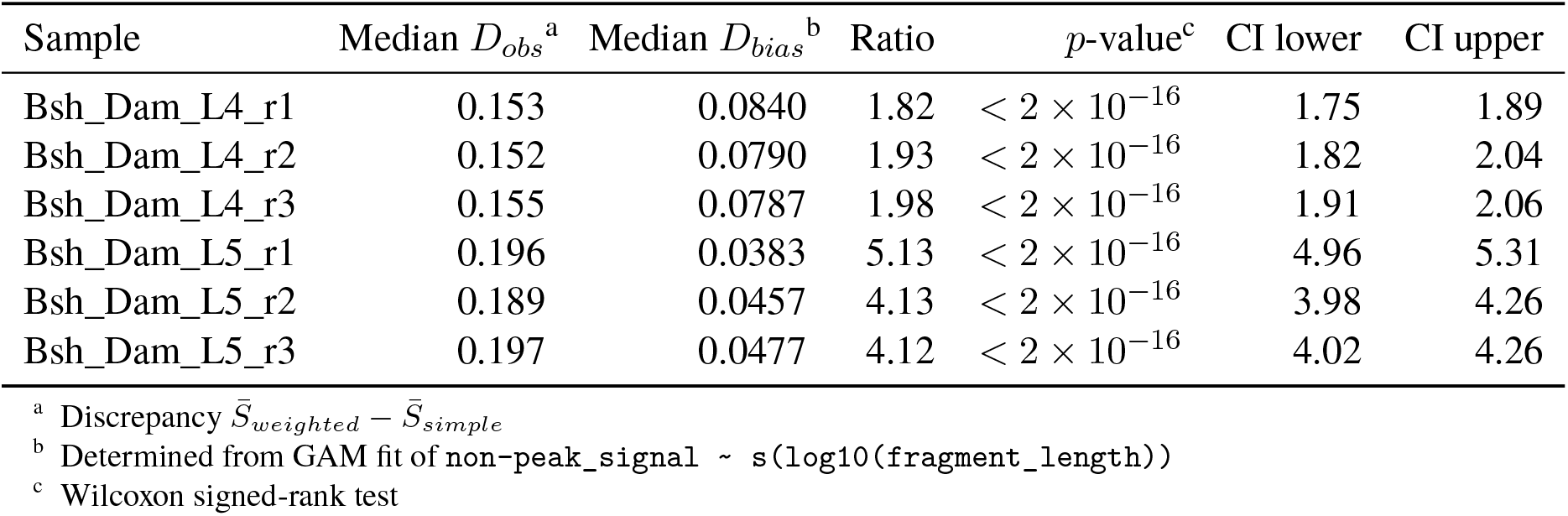
Assessment of fragment-length weighted averaging, Observed vs. Bias-only.

**Table S4:**
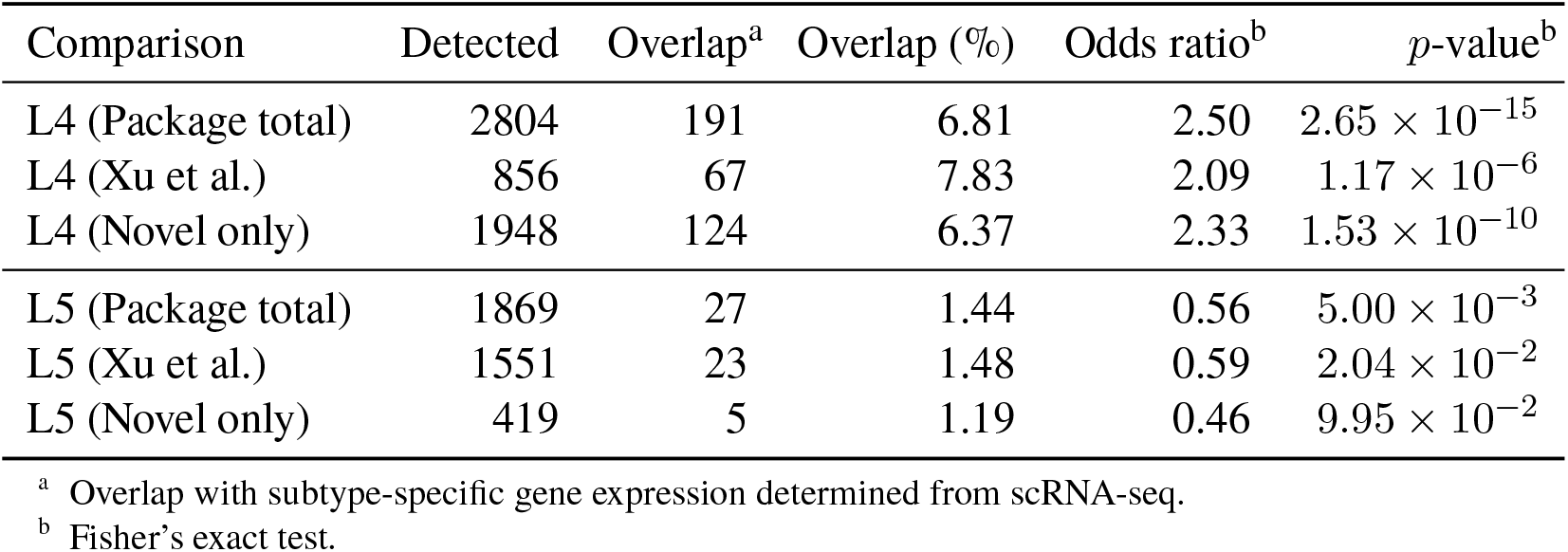
Bsh differential binding overlap with ground-truth scRNA-seq (quantile normalisation)

**Table S5:**
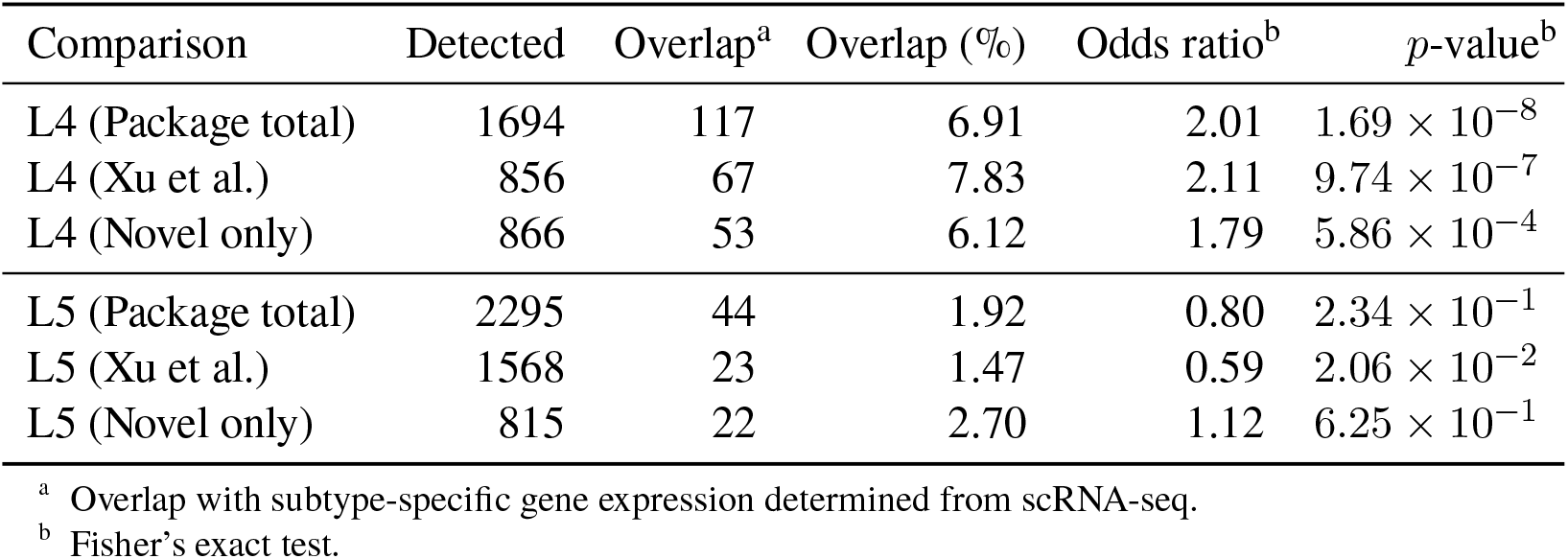
Overlap with ground-truth scRNA-seq (LOESS normalisation)

**Table S6:**
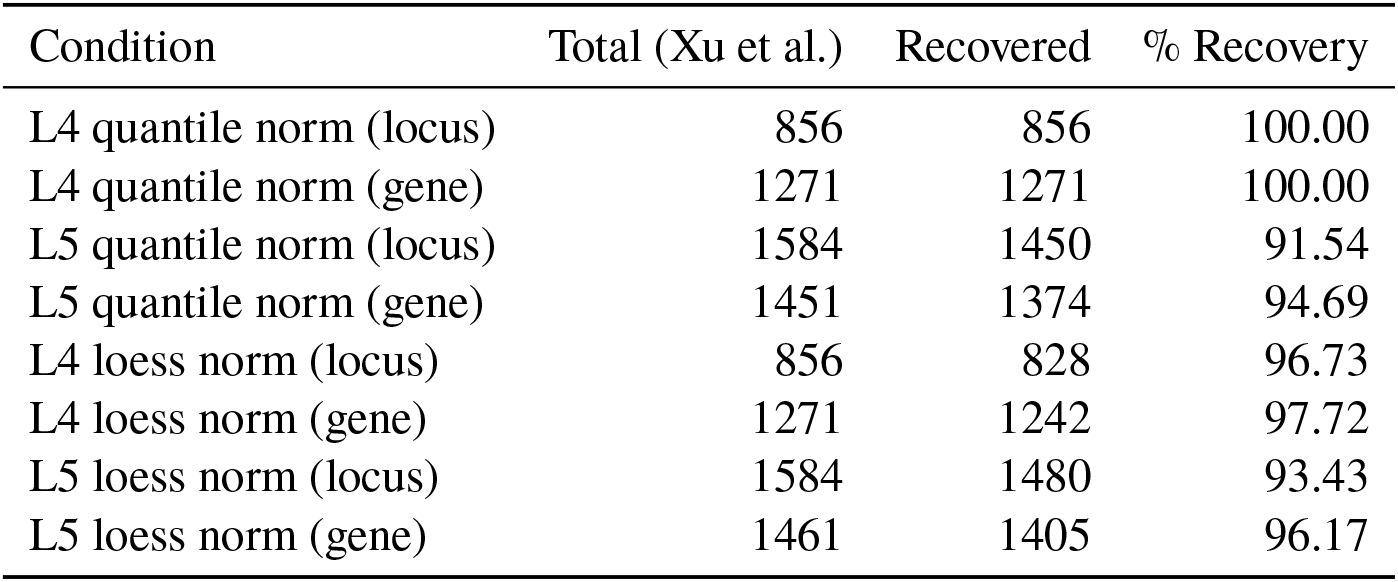
Recovery rates of original loci from Xu et al. across normalisation conditions.

**Table S7:**
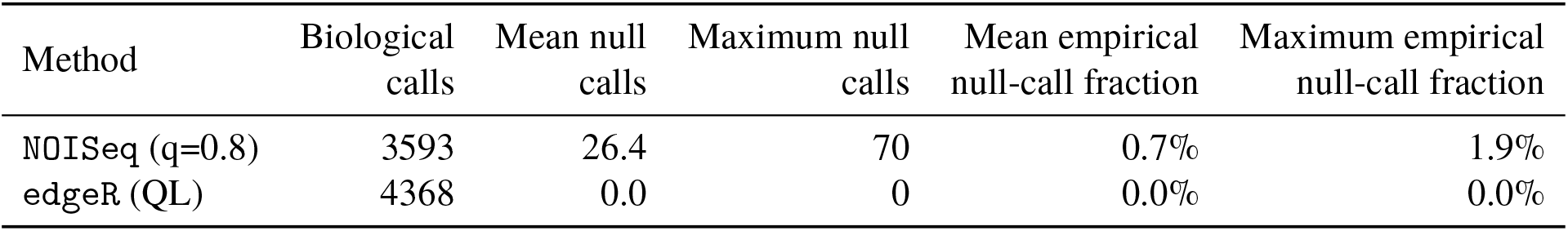
CATaDa permuted null performance by method.

**Table S8:**
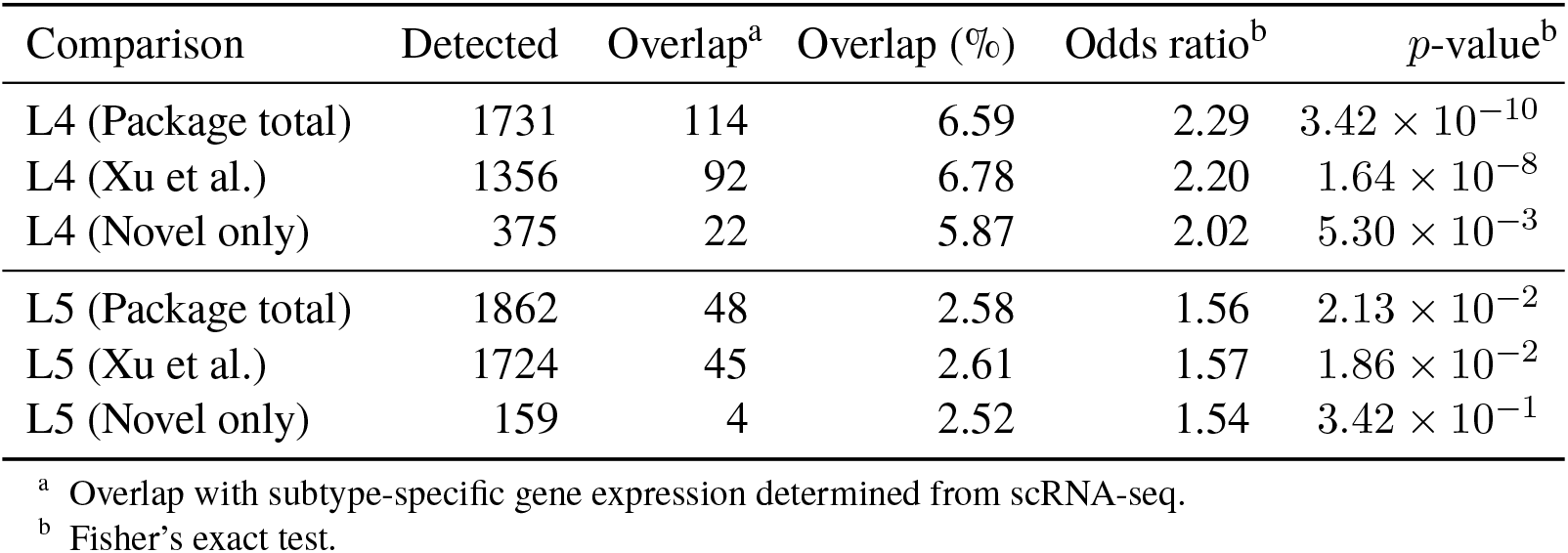
CATaDa differential accessibility overlap with ground-truth scRNA-seq (quantile normalisation)

**Table S9:**
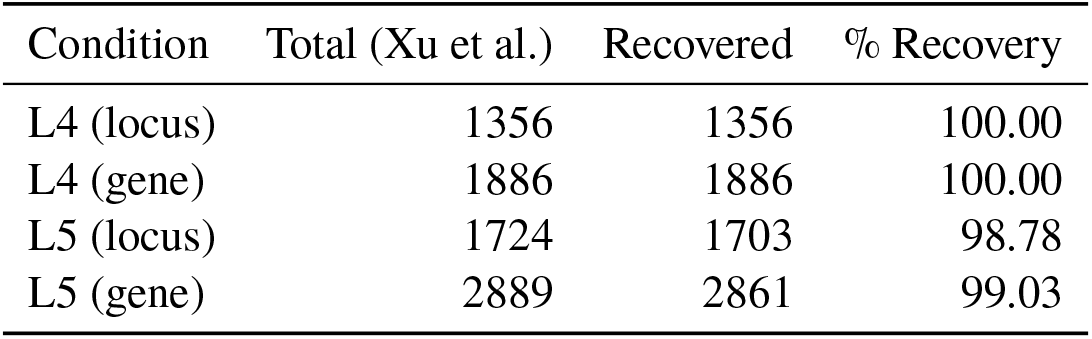
Recovery rates of original CATaDa results from Xu et al.

**Table S10:**
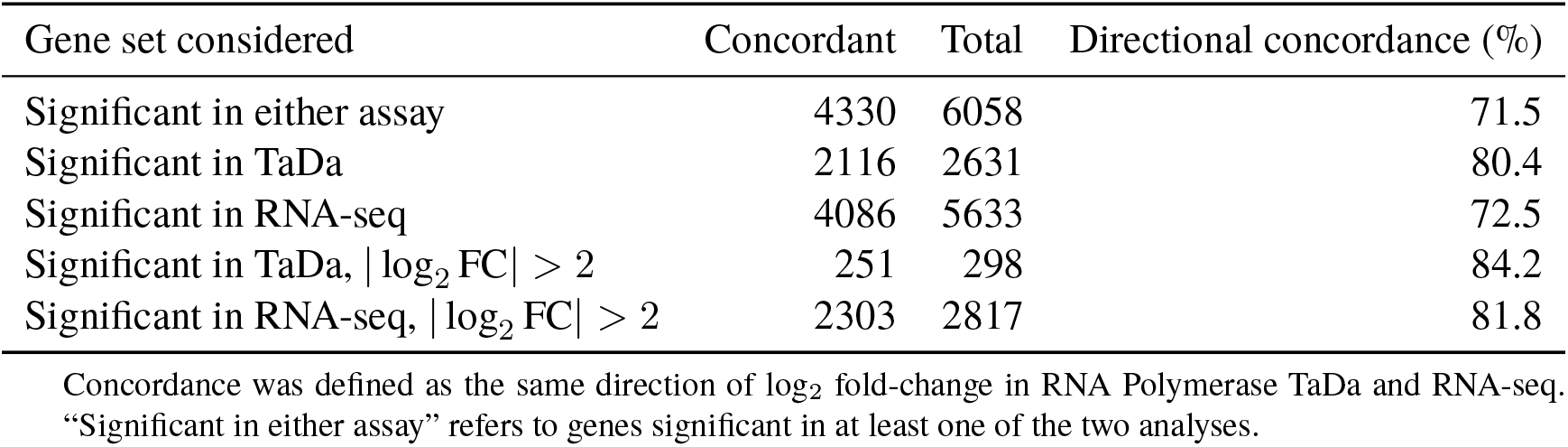
Directional concordance between RNA Polymerase TaDa and bulk RNA-seq differential analysis.

