## Supplementary Material for "damidBind: an R/Bioconductor package for differential DamID analysis and data exploration"

$$\Delta_{obs} = (\bar{B}_w - \bar{B}_s) + (\bar{R}_w - \bar{R}_s).$$

and the corresponding signed discrepancy expected from fragment-length bias alone is

$$\Delta_{bias} = \bar{B}_w - \bar{B}_s$$

Fragment-length bias was modelled for each sample by excluding all fragments overlapping a binding peak, which were considered to contain true signal for which $R_i \gg B(L_i)$. A generalised additive model was then fitted genome-wide to the remaining background fragments:

$$S_i = f(\log_{10}(L_i)) + \epsilon_i$$

where $f$ was estimated using a cubic regression spline. Using this model, the expected length-associated signal $\hat{B}_i$ was predicted for all fragments within tested loci. Because the background set may still contain weak or undetected biological signal, this should provide a conservative upper estimate of fragment-length bias.

For each tested locus containing multiple fragments, the observed absolute discrepancy between the fragment-length-weighted mean and the simple mean was calculated as

$$D_{obs} = |\Delta_{obs}| = |\bar{S}_w - \bar{S}_s|$$

The corresponding absolute discrepancy expected from the fitted fragment-length bias model alone was calculated as

$$D_{bias} = |\Delta_{bias}| = |\bar{\hat{B}}_w - \bar{\hat{B}}_s|$$

These quantities were compared across tested loci and plotted against within-locus fragment-length heterogeneity, measured as the coefficient of variation of fragment widths (CV).

**Algorithm 1:** Sampling points by local isolation

**Input :** $D$: A dataframe containing coordinates
$k$: An integer for both search depth and priority calculation
$r$: A fixed numeric exclusion radius
**Output :** $V_{kept}$: A logical vector indicating points selected for labelling

`# Pre-process, calculate isolation priority and sort`
$X \leftarrow$ Extract $(x, y)$ coordinates from $D$
$X_{scaled} \leftarrow$ Standardise $X$ (centre and scale to unit variance)
$N_{dist} \leftarrow$ For each point in $X_{scaled}$, find distances to its $k$ nearest neighbours
$P \leftarrow$ Calculate the median of $N_{dist}$ for each point $S \leftarrow$ Indices of points sorted by $P$ in descending order

`# Iterative thinning via radius-based exclusion`
Initialise $V_{processed}$ as a boolean vector of size $|D|$ set to *False*
Initialise $V_{kept}$ as a boolean vector of size $|D|$ set to *False*
**foreach** *index* $i \in S$ **do**
    **if** $V_{processed}[i]$ *is False* **then**
        $V_{kept}[i] \leftarrow$ *True*
        $V_{processed}[i] \leftarrow$ *True*
        $R_{idx} \leftarrow$ Indices of all points in $X_{scaled}$ within distance $r$ of $X_{scaled}[i]$
        **foreach** *index* $j \in R_{idx}$ **do**
            $V_{processed}[j] \leftarrow$ *True*
        **end**
    **end**
**end**
**return** $V_{kept}$

**Algorithm 2:** Occupancy FDR: Model training

**Input :** $S_{\text{genome}}$: Binding profile scores
$W_{\text{genome}}$: Fragment widths
$N_{\text{iter}}$: Simulation iterations
$C_{\text{frag}}$: Set of fragment counts
$T_{\text{occ}}$: Set of occupancy thresholds
**Output :** Models $M_{\text{slope}}$, $M_{\text{int}}$, and $M_{\text{MSE}}$

`# Simulate null distribution by sampling fragments`
Initialise matrix $H$ of size $|T_{\text{occ}}| \times |C_{\text{frag}}|$ with zeros
**for** $n \leftarrow 1$ **to** $N_{\text{iter}}$ **do**
  **foreach** $f \in C_{\text{frag}}$ **do**
    Idx $\leftarrow$ Sample $f$ indices from profile with replacement
    $\text{occ}_{\text{rand}} \leftarrow \frac{\sum_{i \in \text{Idx}} S_{\text{genome}}[i] \cdot W_{\text{genome}}[i]}{\sum_{i \in \text{Idx}} W_{\text{genome}}[i]}$ `# Weighted mean of GATC fragments`
    **foreach** $t \in T_{\text{occ}}$ **do**
      **if** $\text{occ}_{\text{rand}} > t$ **then**
        $H[t, f] \leftarrow H[t, f] + 1$
      **end**
    **end**
  **end**
**end**
$P \leftarrow H/N_{\text{iter}}$ `# Empirical probabilities`

`# Tier 1: Log-linear regression for each fragment count`
**foreach** $f \in C_{\text{frag}}$ **do**
  $P_f \leftarrow$ Probability column for $f$; $T_f \leftarrow T_{\text{occ}}$
  Remove indices where $P_f = 0$
  **if** remaining indices $\geq 3$ **then**
    Fit $M_f$: $\log(P_f) \sim T_f$
    Extract coefficients ($\beta_{\text{slope},f}$, $\beta_{\text{int},f}$), standard errors ($\text{SE}_{\text{slope},f}$, $\text{SE}_{\text{int},f}$), and residual variance ($\text{MSE}_f$)
  **end**
**end**

`# Tier 2: Weighted natural spline (NS) regressions`
$f_{\text{log}} \leftarrow \log(f)$
Fit $M_{\text{slope}}$: $\beta_{\text{slope}} \sim \text{NS}(f_{\text{log}}, \text{df} = 3)$ with weights $1/(\text{SE}_{\text{slope}} + \epsilon)$
Fit $M_{\text{int}}$: $\beta_{\text{int}} \sim \text{NS}(f_{\text{log}}, \text{df} = 3)$ with weights $1/(\text{SE}_{\text{int}} + \epsilon)$
Fit $M_{\text{MSE}}$: $\text{MSE} \sim \text{NS}(f, \text{df} = 3)$ `# Unweighted`
**return** $M_{\text{slope}}, M_{\text{int}}, M_{\text{MSE}}$

**Algorithm 3:** Occupancy FDR: $p$-value estimates and replicate integration

**Input :** $G$: Set of genes with observed occupancy and fragment counts per replicate
$C$: Set of test conditions linked to replicates $r$
$M_{\text{slope},r}$, $M_{\text{int},r}$, $M_{\text{MSE},r}$: Pre-trained Tier 2 models for each replicate $r$
Method: Statistical approach (Stouffer or Fisher)
**Output :** $G$ with condition-level adjusted $g.\text{FDR}_c$ values

`# Estimate individual replicate p-values`
**foreach** condition $c \in C$ **do**
  **foreach** replicate $r \in c$ **do**
    **foreach** gene $g \in G$ **do**
      **if** $g$.nfrag $> 0$ **then**
        $\text{pred}_{\text{slope}} \leftarrow$ Predict from $M_{\text{slope},r}$ using $\log(g.\text{nfrag})$
        $\text{pred}_{\text{int}} \leftarrow$ Predict from $M_{\text{int},r}$ using $\log(g.\text{nfrag})$
        $\text{pred}_{\text{MSE}} \leftarrow \max(0,$ Predict from $M_{\text{MSE},r}$ using $g.\text{nfrag})$
        $\log(p) \leftarrow (\text{pred}_{\text{slope}} \times g.\text{occ}) + \text{pred}_{\text{int}}$
        $p_{g,r} \leftarrow \exp(\log(p) + \text{pred}_{\text{MSE}}/2)$ `# Jensen's correction`
        $p_{g,r} \leftarrow \text{clamp}(p_{g,r}, \text{min} = 10^{-16}, \text{max} = 1.0)$
      **end**
      **else**
        $p_{g,r} \leftarrow 1.0$
      **end**
    **end**
  **end**

  `# Aggregate p-values across replicates per condition`
  **foreach** gene $g \in G$ **do**
    $k \leftarrow$ Count of replicates for $g$ where $p_{g,r} < 1.0$
    **if** $k = 1$ **then**
      $P_{\text{combined},g} \leftarrow$ the single non-1.0 p-value
    **end**
    **else if** Method is Stouffer **then**
      **foreach** $r \in c$ **do**
        $z_r \leftarrow \Phi^{-1}(1 - p_{g,r})$ `# Inverse Normal CDF`
      **end**
      $Z_{\text{meta}} \leftarrow \frac{\sum z_r}{\sqrt{k}}$ `# Unweighted combination`
      $P_{\text{combined},g} \leftarrow 1 - \Phi(Z_{\text{meta}})$ `# Back-transform`
    **end**
    **else if** Method is Fisher **then**
      $X^2 \leftarrow -2\sum_{r=1}^{k} \log(p_{g,r})$ `# Fisher statistic`
      $P_{\text{combined},g} \leftarrow P(\chi^2_{2k} > X^2)$ `# Chi-squared lookup`
    **end**
  **end**

  `# FDR correction at condition level`
  $G.\text{FDR}_c \leftarrow \text{Benjamini-Hochberg}(P_{\text{combined}})$
  **return** $G$ with assigned $G.\text{FDR}_c$
**end**

### Supplementary figures

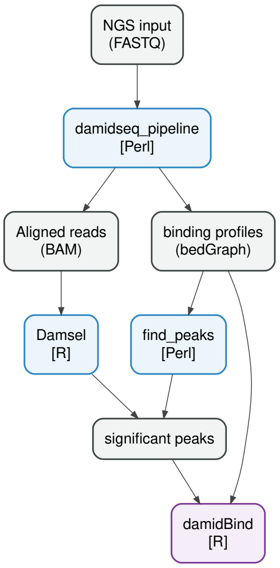


**Figure S1:** Suggested workflow options for DamID-seq data processing, from raw sequencing reads to `damidBind`.

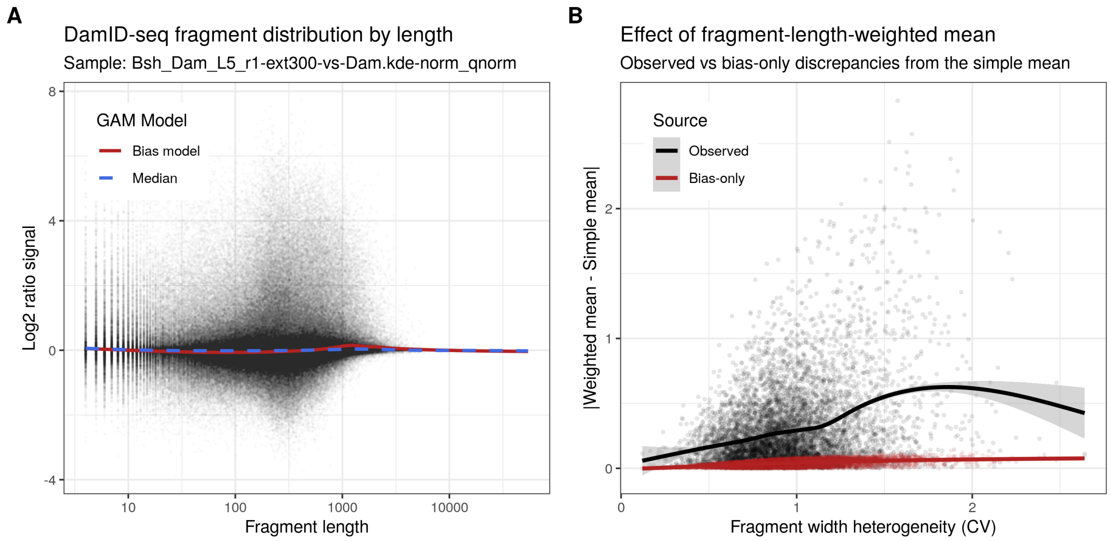


**Figure S2:** Discrepancies between fragment-length-weighted averaging and simple averaging are not driven by systematic bias. (A) Plots of the underlying fragment length bias over all non-peak GATC fragments, used to fit a null model of systematic fragment-length signal bias (red); a fit of the signal median is also shown. (B) Discrepancy between simple and fragment-length-weighted means, both from the data (Observed, black) and as predicted using the bias-only model (Bias-only, red).

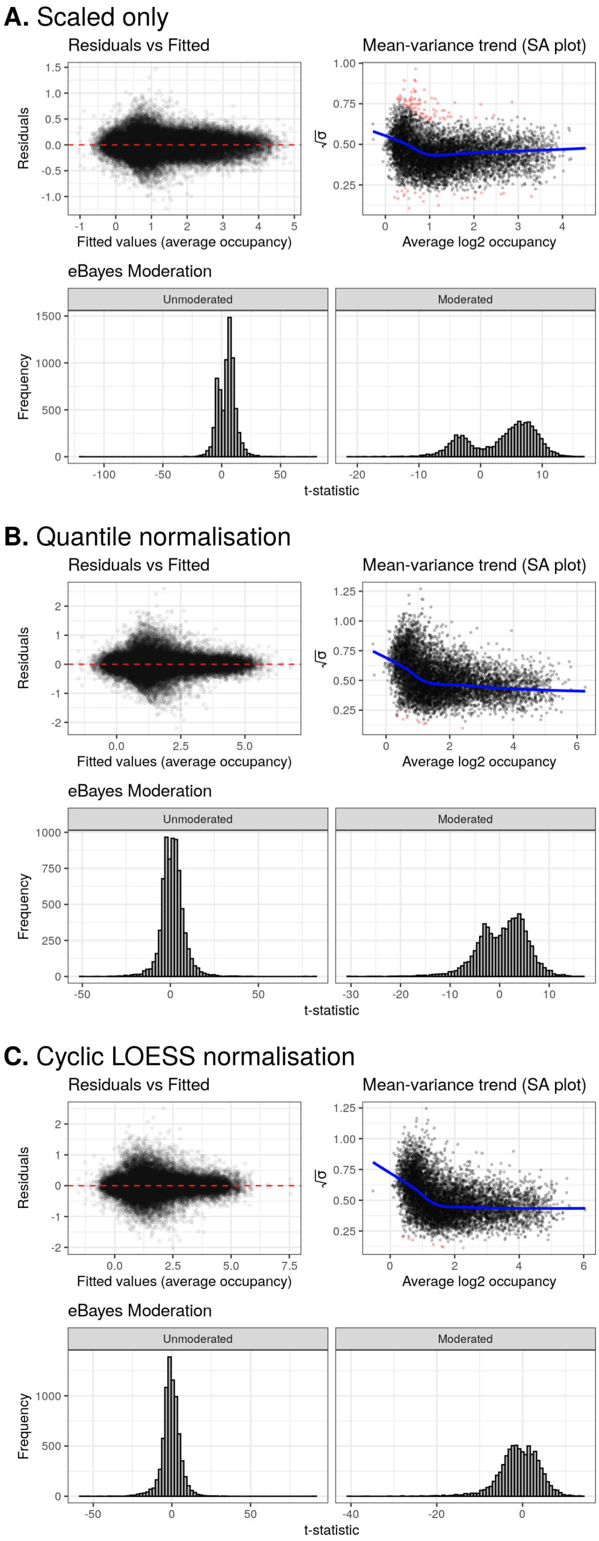


**Figure S3:** Diagnostic plots for `limma`'s eBayes moderation generated by `damidBind` using Bsh L4/L5 binding dataset with only zero-preserving scaling, with quantile normalisation, and with cyclic LOESS normalisation. Plots showing the heteroscedasticity of the data, the SA plot showing the model fit to the mean-variance of the data (outliers coloured red; fit line in blue), and the effect of eBayes moderation on t-statistic shrinkage.

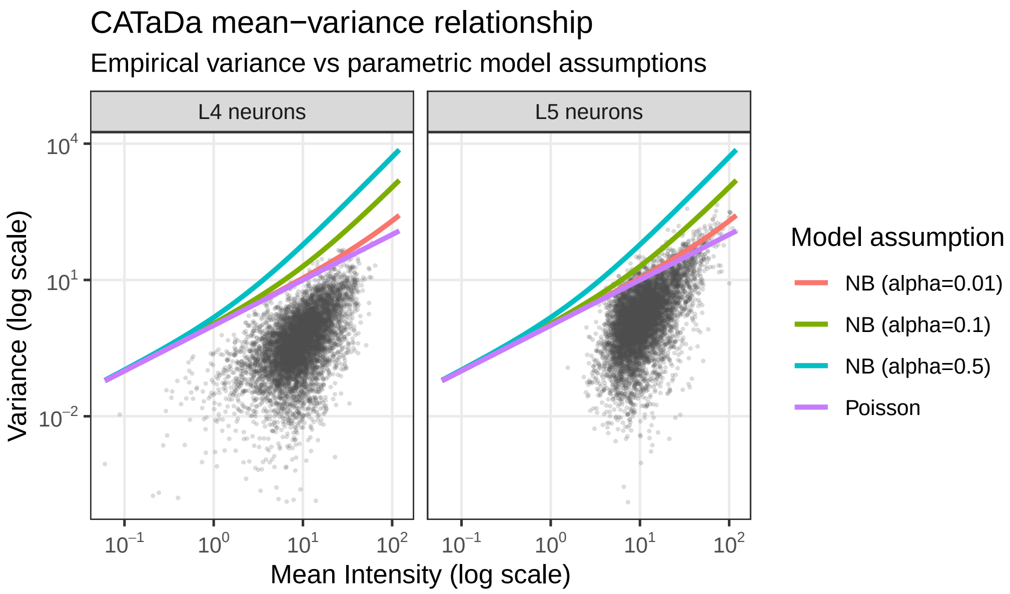


**Figure S4:** Plots of mean vs variance relationship of CATaDa peak data. Data for L4 and L5 neuronal subtypes (from (Xu *et al.*, 2024)) are shown, alongside Poisson (Var $= \mu$) and Negative Binomial (Var $= \mu + \alpha\mu^2$) responses; points represent individual loci. Input data was loaded with `pre_scale=FALSE`, `norm_method="none"` (i.e. no normalisation was applied to the dataset). In both cases, the data sit below the model lines and are under-dispersed, violating NB and Poisson assumptions.

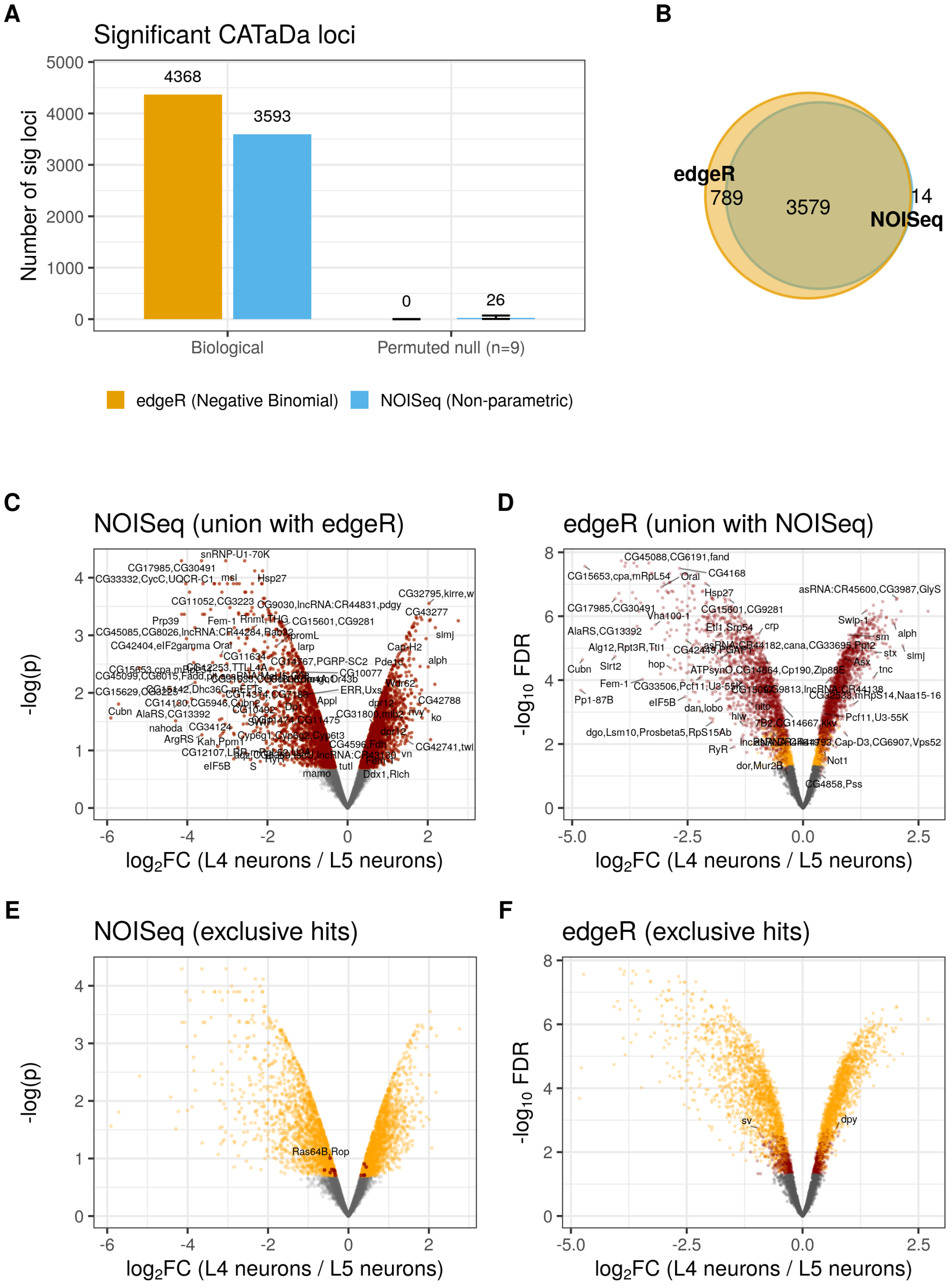


**Figure S5:** Comparison of `NOISeq` and `edgeR` performance on CATaDa data. (A) Significant hits generated by both methods, for both the biological lamina neuron data from Xu *et al.* (2024) and a permuted null dataset which should yield no hits, to assess FDR performance. (B) Venn diagram of overlapping significant loci called by the two methods on the lamina dataset. (C and D) Significant `NOISeq` and `edgeR` loci, coloured by set overlap in red, other significant loci in orange. (E and F) Significant `NOISeq` and `edgeR` loci, coloured by set uniqueness in red, other significant loci in orange.

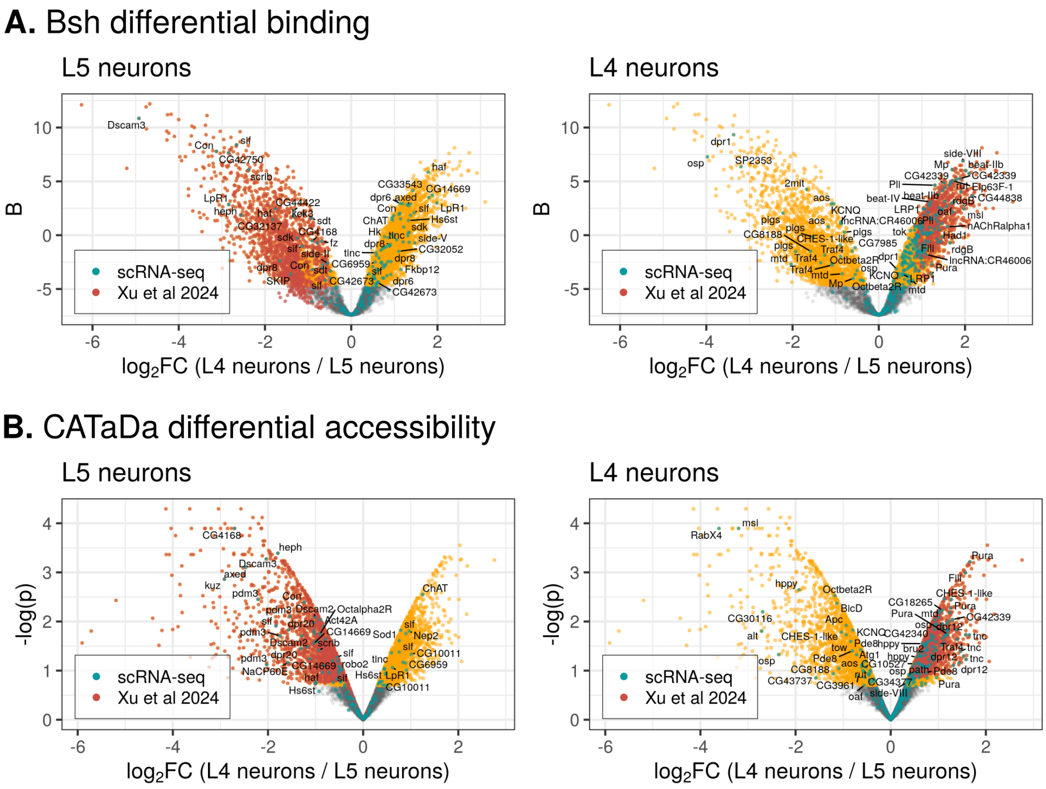


**Figure S6:** Comparison of `damidBind`'s performance against previously published analysis from Xu *et al.* (2024). (A) Bsh TF differential binding; (B) CATaDa differential accessibility. In all plots, significant loci identified by `damidBind` are coloured orange; significant loci identified by Xu *et al.* (2024) are coloured red; scRNA-seq gene expression markers uniquely expressed in each lineage (data from (Xu *et al.*, 2024)) are coloured cyan. See also Table S4. Results using quantile normalisation are shown.

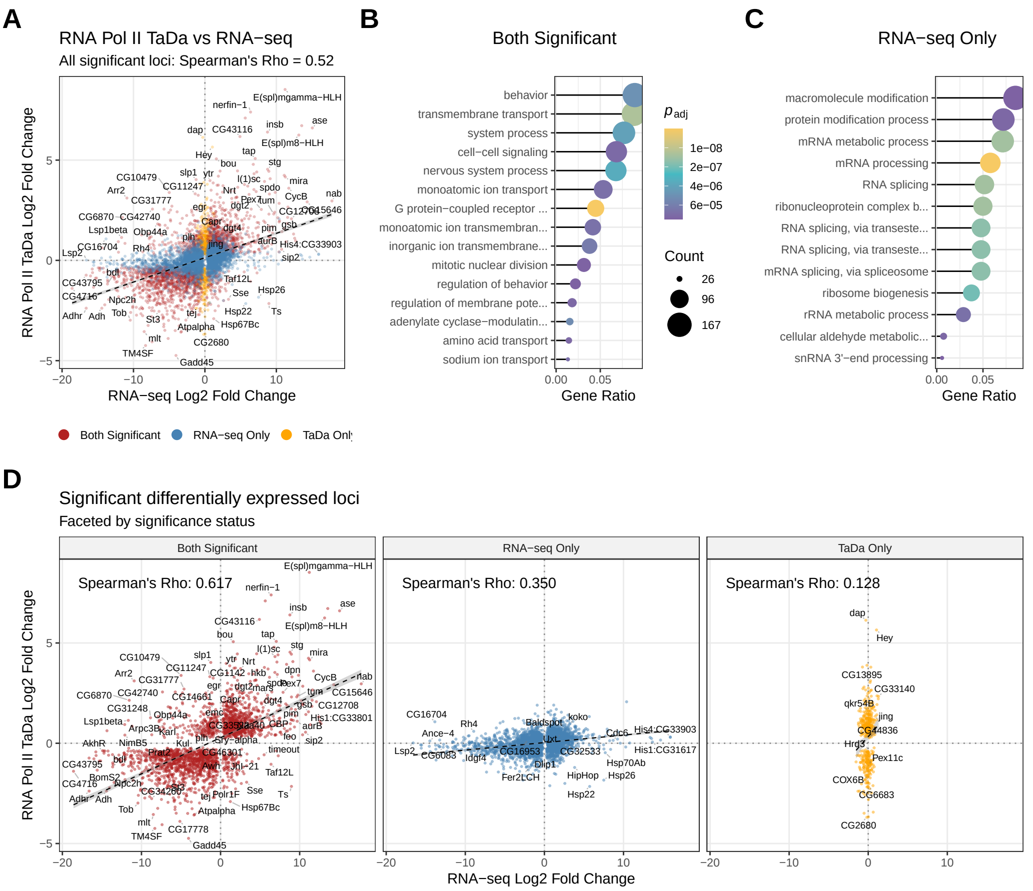


**Figure S7:** Differential expression compared between TaDa and RNA-seq. In both cases, third instar larval neural stem cells were compared to adult neurons. (A) All significant loci compared; (B) faceted by significance status; (C) Gene Ontology (GO) enrichment terms by significance class (no GO terms were significantly enriched for the TaDa-only significance group). See Supplementary Methods for details.

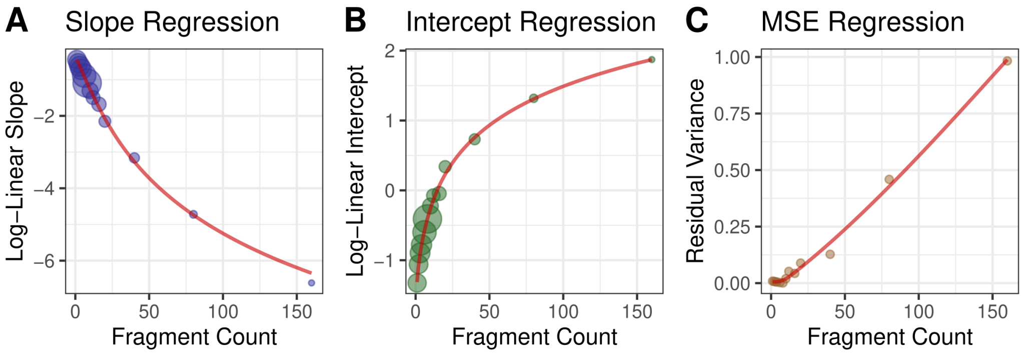


**Figure S8:** FDR modelling for RNA Polymerase occupancy data. The plots show the Tier 2 regression fits and Tier 1 model heteroscedasticity, for a typical DamID RNA Polymerase dataset. Both the Tier 2 slope (A) and intercept (B) regressions use weighted natural spline fits to fit the underlying data; circle size is proportional to weight. The model mean squared error (MSE) increases with fragment count (C) and is also fitted with an unweighted natural spline to apply the appropriate model bias correction upon back-transformation. Data illustrated is derived from RNA Polymerase (RpII215) occupancy data from GSE77860.

### Supplementary tables

**Table S1:** Summary of DamID-seq-related bioinformatics software

| **Package** | **Function** | **Outputs** |
|---|---|---|
| `damidseq_pipeline` | NGS sequencing alignment, Dam-fusion/Dam-only signal normalisation, and $\log_2$ ratio generation, or RPM occupancy counts in the case of CATaDa. | bedGraph profiles and BAM alignments |
| `Damsel` | Identifies significant peaks from individual replicates of Dam-fusion/Dam-only DamID-seq data, using BAM inputs. | `dm_results` object; `peaks(dm_results)` provides the peaks as a `GenomicRanges` object |
| `find_peaks` | Per-replicate peak caller on $\log_2$ ratio datasets. Finds contiguous high-signal regions that pass an FDR threshold. | BED or GFF peak files |
| `damidBind` | Performs differential binding, expression, or accessibility analysis on input profiles generated by `damidseq_pipeline`, along with peaks generated via `Damsel` or `find_peaks`. | `DamIDResults` object |

**Table S2:** Benchmark results for `damidBind` workflows using serial and multicore execution. Significantly shorter execution times result from parallel processing, although the benefits of using more than 15 cores is limited.

| Workflow | Execution backend[a] | Median | Min | Total time | $n_{\text{itr}}$ |
|---|---|---|---|---|---|
| `load_data_genes()` | serial | 15.11 min | 15.08 min | 1.26 h | 5 |
| `load_data_genes()` | multicore_15 | 1.80 min | 1.78 min | 9.23 min | 5 |
| `load_data_genes()` | multicore_30 | 1.70 min | 1.67 min | 8.47 min | 5 |
| `load_data_peaks()` | serial | 1.16 min | 1.15 min | 5.82 min | 5 |
| `load_data_peaks()` | multicore_15 | 13.99 s | 13.21 s | 1.15 min | 5 |
| `load_data_peaks()` | multicore_30 | 12.06 s | 11.89 s | 1.06 min | 5 |

[a] Execution backend corresponds to: `BiocParallel::SerialParam()`, `BiocParallel::MulticoreParam(workers = 15)`, and `BiocParallel::MulticoreParam(workers = 30)` respectively.

**Table S3:** Assessment of fragment-length weighted averaging, Observed vs. Bias-only.

| Sample | Median $D_{obs}$[a] | Median $D_{bias}$[b] | Ratio | $p$-value[c] | CI lower | CI upper |
|---|---|---|---|---|---|---|
| Bsh_Dam_L4_r1 | 0.153 | 0.0840 | 1.82 | $< 2 \times 10^{-16}$ | 1.75 | 1.89 |
| Bsh_Dam_L4_r2 | 0.152 | 0.0790 | 1.93 | $< 2 \times 10^{-16}$ | 1.82 | 2.04 |
| Bsh_Dam_L4_r3 | 0.155 | 0.0787 | 1.98 | $< 2 \times 10^{-16}$ | 1.91 | 2.06 |
| Bsh_Dam_L5_r1 | 0.196 | 0.0383 | 5.13 | $< 2 \times 10^{-16}$ | 4.96 | 5.31 |
| Bsh_Dam_L5_r2 | 0.189 | 0.0457 | 4.13 | $< 2 \times 10^{-16}$ | 3.98 | 4.26 |
| Bsh_Dam_L5_r3 | 0.197 | 0.0477 | 4.12 | $< 2 \times 10^{-16}$ | 4.02 | 4.26 |

[a] Discrepancy $\bar{S}_{weighted} - \bar{S}_{simple}$
[b] Determined from GAM fit of `non-peak_signal ~ s(log10(fragment_length))`
[c] Wilcoxon signed-rank test

**Table S4:** Bsh differential binding overlap with ground-truth scRNA-seq (quantile normalisation)

| Comparison | Detected | Overlap[a] | Overlap (%) | Odds ratio[b] | $p$-value[b] |
|---|---|---|---|---|---|
| L4 (Package total) | 2804 | 191 | 6.81 | 2.50 | $2.65 \times 10^{-15}$ |
| L4 (Xu et al.) | 856 | 67 | 7.83 | 2.09 | $1.17 \times 10^{-6}$ |
| L4 (Novel only) | 1948 | 124 | 6.37 | 2.33 | $1.53 \times 10^{-10}$ |
| L5 (Package total) | 1869 | 27 | 1.44 | 0.56 | $5.00 \times 10^{-3}$ |
| L5 (Xu et al.) | 1551 | 23 | 1.48 | 0.59 | $2.04 \times 10^{-2}$ |
| L5 (Novel only) | 419 | 5 | 1.19 | 0.46 | $9.95 \times 10^{-2}$ |

[a] Overlap with subtype-specific gene expression determined from scRNA-seq.
[b] Fisher's exact test.

**Table S5:** Overlap with ground-truth scRNA-seq (LOESS normalisation)

| Comparison | Detected | Overlap[a] | Overlap (%) | Odds ratio[b] | $p$-value[b] |
|---|---|---|---|---|---|
| L4 (Package total) | 1694 | 117 | 6.91 | 2.01 | $1.69 \times 10^{-8}$ |
| L4 (Xu et al.) | 856 | 67 | 7.83 | 2.11 | $9.74 \times 10^{-7}$ |
| L4 (Novel only) | 866 | 53 | 6.12 | 1.79 | $5.86 \times 10^{-4}$ |
| L5 (Package total) | 2295 | 44 | 1.92 | 0.80 | $2.34 \times 10^{-1}$ |
| L5 (Xu et al.) | 1568 | 23 | 1.47 | 0.59 | $2.06 \times 10^{-2}$ |
| L5 (Novel only) | 815 | 22 | 2.70 | 1.12 | $6.25 \times 10^{-1}$ |

[a] Overlap with subtype-specific gene expression determined from scRNA-seq.
[b] Fisher's exact test.

**Table S6:** Recovery rates of original loci from Xu et al. across normalisation conditions

| Condition | Total (Xu et al.) | Recovered | % Recovery |
|---|---|---|---|
| L4 quantile norm (locus) | 856 | 856 | 100.00 |
| L4 quantile norm (gene) | 1271 | 1271 | 100.00 |
| L5 quantile norm (locus) | 1584 | 1450 | 91.54 |
| L5 quantile norm (gene) | 1451 | 1374 | 94.69 |
| L4 loess norm (locus) | 856 | 828 | 96.73 |
| L4 loess norm (gene) | 1271 | 1242 | 97.72 |
| L5 loess norm (locus) | 1584 | 1480 | 93.43 |
| L5 loess norm (gene) | 1461 | 1405 | 96.17 |

**Table S7:** CATaDa permuted null performance by method.

| Method | Biological calls | Mean null calls | Maximum null calls | Mean empirical null-call fraction | Maximum empirical null-call fraction |
|---|---|---|---|---|---|
| `NOISeq` (q=0.8) | 3593 | 26.4 | 70 | 0.7% | 1.9% |
| `edgeR` (QL) | 4368 | 0.0 | 0 | 0.0% | 0.0% |

**Table S8:** CATaDa differential accessibility overlap with ground-truth scRNA-seq (quantile normalisation)

| Comparison | Detected | Overlap[a] | Overlap (%) | Odds ratio[b] | $p$-value[b] |
|---|---|---|---|---|---|
| L4 (Package total) | 1731 | 114 | 6.59 | 2.29 | $3.42 \times 10^{-10}$ |
| L4 (Xu et al.) | 1356 | 92 | 6.78 | 2.20 | $1.64 \times 10^{-8}$ |
| L4 (Novel only) | 375 | 22 | 5.87 | 2.02 | $5.30 \times 10^{-3}$ |
| L5 (Package total) | 1862 | 48 | 2.58 | 1.56 | $2.13 \times 10^{-2}$ |
| L5 (Xu et al.) | 1724 | 45 | 2.61 | 1.57 | $1.86 \times 10^{-2}$ |
| L5 (Novel only) | 159 | 4 | 2.52 | 1.54 | $3.42 \times 10^{-1}$ |

[a] Overlap with subtype-specific gene expression determined from scRNA-seq.
[b] Fisher's exact test.

**Table S9:** Recovery rates of original CATaDa results from Xu et al.

| Condition | Total (Xu et al.) | Recovered | % Recovery |
|---|---|---|---|
| L4 (locus) | 1356 | 1356 | 100.00 |
| L4 (gene) | 1886 | 1886 | 100.00 |
| L5 (locus) | 1724 | 1703 | 98.78 |
| L5 (gene) | 2889 | 2861 | 99.03 |

**Table S10:** Directional concordance between RNA Polymerase TaDa and bulk RNA-seq differential analysis

| Gene set considered | Concordant | Total | Directional concordance (%) |
|---|---|---|---|
| Significant in either assay | 4330 | 6058 | 71.5 |
| Significant in TaDa | 2116 | 2631 | 80.4 |
| Significant in RNA-seq | 4086 | 5633 | 72.5 |
| Significant in TaDa, $\lvert \log_2 \mathrm{FC} \rvert > 2$ | 251 | 298 | 84.2 |
| Significant in RNA-seq, $\lvert \log_2 \mathrm{FC} \rvert > 2$ | 2303 | 2817 | 81.8 |

Concordance was defined as the same direction of $\log_2$ fold-change in RNA Polymerase TaDa and RNA-seq. "Significant in either assay" refers to genes significant in at least one of the two analyses.
